# *De novo* minibinders targeting RH5 achieve nanomolar inhibition of blood-stage malaria replication

**DOI:** 10.64898/2026.09.28.755224

**Authors:** Jason Hsiao, Melina Shamshoum, Abhiram Chalamalasetty, Jordan R. Barrett, Ricardo Almada-Monter, Elizabeth Tabornal, Emily Wilks, Yi-Zong Lee, Nathan Beutler, Amanda R. Lukens, Priyan Kapoor, Daisy Chen, Justice Ndihokubwayo, Barnabas G. Williams, Kirsty McHugh, David Pulido, Irina Mendes de Sousa, Elizabeth Komives, Emília V Noormahomed, Ian A. Wilson, Dyann F. Wirth, Thomas Rogers, Neville Bethel, Simon J. Draper, Andrew Ward, Elizabeth A. Winzeler, Adrian Jinich

## Abstract

Malaria kills ∼600,000 people annually despite recently deployed vaccines. The parasite’s blood-stage invasion of erythrocytes, driven by the essential, non-redundant interaction between *Plasmodium falciparum* RH5 and erythrocyte basigin, remains a leading therapeutic target. Here, we designed *de novo* mini-proteins (“minibinders”) to occlude this interface. From ∼1,500 computationally generated designs, we synthesized 18 candidates ranging from 9.3 to 16.9 kDa which were easily expressed in *E. coli* as soluble, well-folded proteins. Of these, 7 minibinders blocked blood-stage replication of *P. falciparum* asexual stage parasites with greater potency than ≈300nM IC_50_. Three potent candidate minibinders, mb-5, mb-7, and mb-21, were confirmed to bind RH5 via SPR and BLI and block the merozoite-to-ring transition in stage-synchronized parasite progression assays, with CryoEM of minibinder-RH5 complex confirming the predicted binding at RH5-basigin interface. Although pharmacological challenges remain, our data demonstrate that potent inhibitors of parasite growth, rivaling existing antimalarials, can be designed rather than discovered via extensive screening. Our data highlight the potential for artificial intelligence to change the development landscape of new antimalarial therapies.

## Main

Malaria remains a leading cause of morbidity and mortality worldwide, with *Plasmodium falciparum* responsible for most severe cases and deaths. In 2024, malaria caused an estimated 282 million cases and 610,000 deaths (1). Although the recently deployed RTS,S/AS01 (2021) and R21/Matrix-M (2023) marked the first malaria vaccines to reach broad use, antibody titers wane rapidly. Drug- and insecticide-resistance also continue to erode existing control strategies. Highly effective antimalarial therapeutics and prophylactics against essential parasite functions remain a critical, yet unmet need.

Several of our current antimalarial therapies were discovered by extracting the active ingredient from folk remedies with fever-reducing properties and optimizing drug-like properties (artemisinin, chloroquine). More recently, the field has used high-throughput screening to find therapeutic candidates. The starting point for Ganaplacide, the most clinically advanced antimalarial with novel mechanism of action, was discovered by testing millions of drug candidates for their ability to block parasite growth using miniaturized parasite assays (2). Likewise, monoclonal antibodies that are currently in clinical trials, albeit against pre-erythrocytic sporozoites (3, 4), were discovered by first screening thousands of reverse-engineered monoclonal antibodies produced by vaccinated individuals for binding and inhibition of parasite invasion. While such efforts have yielded highly potent monoclonal antibodies (5-9), this discovery process is laborious and inefficient.

*De novo* protein design has rapidly matured to the point where computationally designed minibinders can achieve antibody-like binding affinities, while offering substantial advantages over antibodies: small size, high thermostability, absence of disulfide bonds, and amenability to low-cost bacterial expression (10-13). Recent advances in structure prediction (AlphaFold2/3, Boltz-2) and sequence design (ProteinMPNN) have dramatically improved design success rates (14-17), with some even achieving picomolar potencies in infectious disease contexts (18).

To test the hypothesis that *de novo* protein design could be used to find candidates that block malaria parasite development, we selected reticulocyte-binding protein homolog 5 (RH5) as a well-validated target. The blood-stage of the *P. falciparum* life cycle, during which merozoites invade and multiply within erythrocytes, is responsible for all clinical manifestations of malaria (***Figure 1a***). Invasion requires a complex choreography of molecular interactions, among which the RH5–CyRPA–RIPR (RCR) complex, anchored by PTRAMP and CSS (“PCRCR-complex”), plays an indispensable role (***Figure 1b***) (19-21). Unlike other invasion ligands, RH5 has no functional redundancy: genetic knockout of RH5 is lethal to the parasite. RH5 mediates invasion through its essential interaction with basigin (CD147/BSG) (22), a broadly expressed glycoprotein on the erythrocyte surface, and it is highly conserved across *P. falciparum* strains (23), minimizing the risk of resistance development through epitope variation. Although it has not been targeted by small molecules, anti-RH5 monoclonal antibodies (mAb) have demonstrated potent neutralizing activity in growth-inhibition assays (GIA) (7), and remains one of the most advanced blood-stage vaccine and antibody targets in clinical development (24). Critically, the most potent neutralizing antibodies act by either directly overlapping the basigin-binding site (e.g., R5.004) or binding adjacent regions that sterically hinder the interaction (e.g., 9AD4, R5.016, R5.034) (5, 6, 9). Past structural studies utilizing these potently neutralizing antibodies provide excellent structural information and reveal a clear design strategy: direct occlusion of the RH5-basigin binding interface using small, rigid minibinders computationally biased to engage multiple RH5 hotspot residues simultaneously.

**Figure 1.**
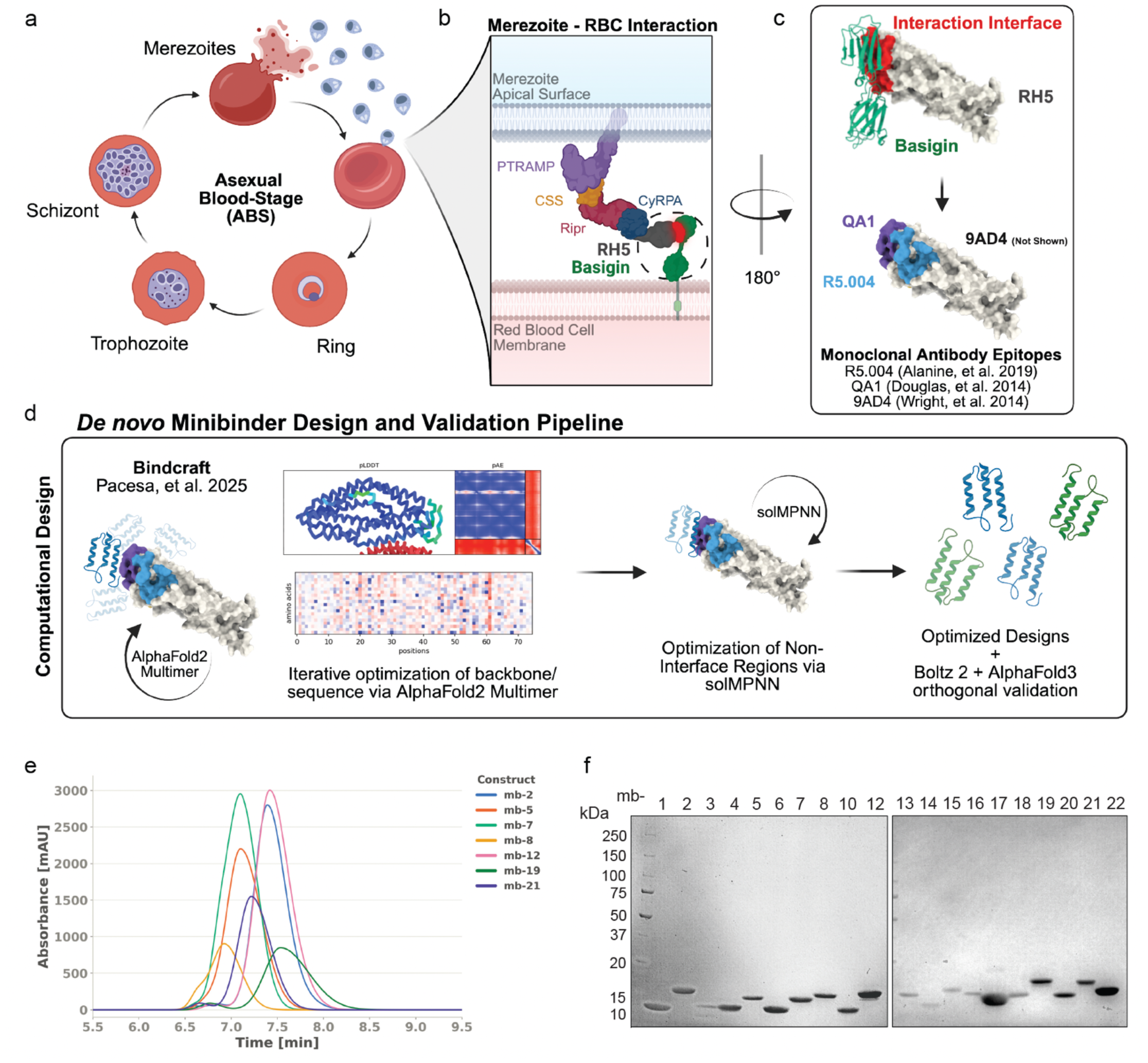
The RH5-Basigin invasion interface and a *de novo* minibinder inhibitor design strategy. (**a**) Asexual blood-stage life cycle of *Plasmodium falciparu*m. (**b**) Schematic of RH5 (grey) on the merozoite surface engaging basigin (green, CD147) on the erythrocyte membrane. (**c**) Three RH5 residues chosen as design constraints from the basigin-contacting residues and epitopes of known neutralizing anti-RH5 monoclonal antibodies (R5.004, 9AD4, QA1). (**d**) *De novo* design and validation pipeline. (**e**) Size-exclusion HPLC traces of seven representative monoclonal minibinder constructs used for downstream assays. (**f**) SDS-PAGE analysis of 20 recombinantly expressed minibinders (mb1 - mb22, labeled by number), each resolving as a single band near the predicted molecular weights (∼9-17 kDa).

### *De novo* design of minibinders targeting the RH5–basigin interface

To test whether *de novo* protein design could generate inhibitors of RH5-dependent erythrocyte invasion, we designed and computationally validated a panel of minibinders against the *Pf*RH5– basigin interface. Using the BindCraft pipeline (12), we generated ∼1,500 *de novo* minibinder designs targeting the basigin-binding interface of *Pf*RH5 (PDB: 4WAT) (25). BindCraft uses iterative backpropagation through the AlphaFold2 multimer network to simultaneously hallucinate and optimize *de novo* protein sequences that bind a target protein with high predicted affinity and specificity. Specifically, designs were constrained to simultaneously contact three hotspot residues (449, 358, 201) located near the kite-tip of RH5 within the basigin-binding site. These design constraints resulted from two convergent lines of evidence: structural analysis of the RH5–basigin contact interface (PDB: 4U0Q) (9) and the epitopes of neutralizing anti-RH5 monoclonal antibodies (R5.004, 9AD4, QA1) (5, 7, 9) (***Figure 1c***). Sequences were initialized randomly, iteratively optimized using a composite loss function (structural confidence, interface quality, predicted alignment error, residue contacts, compactness) within BindCraft’s modified AlphaFold2 framework, and refined with SolMPNN. Final designs were filtered against stringent criteria (i-PAE < 0.35, pLDDT > 0.86, Rosetta ΔG < −45 kcal/mol, ΔG/ΔSASA < −2.0 kcal/mol/Å^2^; design length 75–175 residues; helicity-favored; cysteine-free). Top candidates were independently validated using AlphaFold3 (14) and Boltz-2 (17). Validation metrics confirmed high-confidence binding interfaces (ipTM 0.85–0.93) with no detected steric clashes (***Table S1***).

### Computationally designed minibinders are well-expressed and bind RH5

Having generated a panel of high-confidence designs *in silico*, we next asked whether they could be produced as stable, folded proteins and whether they engage RH5 with the affinity predicted by the design models. Strikingly, 100% of our 19 computationally filtered scaffolds (77–166 aa, median 114 aa) were successfully expressed in *E. coli* and purified to homogeneity using standard affinity and size-exclusion chromatography (SEC). SEC profiles showed that most designs adopted a predominantly monomeric state; where higher-order species were present, the monomeric fraction was resolved and pooled for downstream characterization (***Figure 1e, S1, S2***).

To identify which designs engage RH5 in solution, we screened the purified panel by biolayer interferometry (BLI), immobilizing recombinant RH5 and monitoring association and dissociation of minibinders. BLI screening showed a continuum of RH5 binding response with mb-21 possessing the greatest binding response (***Figure 2a***). Because BLI lacks bulk flow and its kinetics are mass-transport-limited, we used BLI only as a qualitative binding readout and then validated and measured binding kinetics of a subset of ten minibinders by surface plasmon resonance (SPR) (***Figure 2b, S3, Table S2***). Three leads (mb-5, mb-7, and mb-21), which were characterized in subsequent parasite assays, bound RH5 with apparent K_D_ values in the low-micromolar range (mb-21, 4.8 µM; mb-5, 11.2 µM; mb-7, 11.8 µM) (***Figure 2b, S3, Table S2***), with mb-21 combining the highest apparent affinity with the slowest off-rate (apparent k_off_ ≈ 1.0 × 10^−2^ s^−1^). Specifically, mb-21-RH5 complex persisted for ∼100s on average with a dissociation half-life of ∼70s, roughly 4 to 7-fold longer than all other binders characterized. SPR results were fully concordant with BLI response. Every minibinder that bound RH5 by SPR produced a positive BLI response, while all four SPR non-binders (mb-8, mb-13, mb-17, mb18) gave BLI responses at or below baseline (***Figure 2a, S3***).

**Figure 2.**
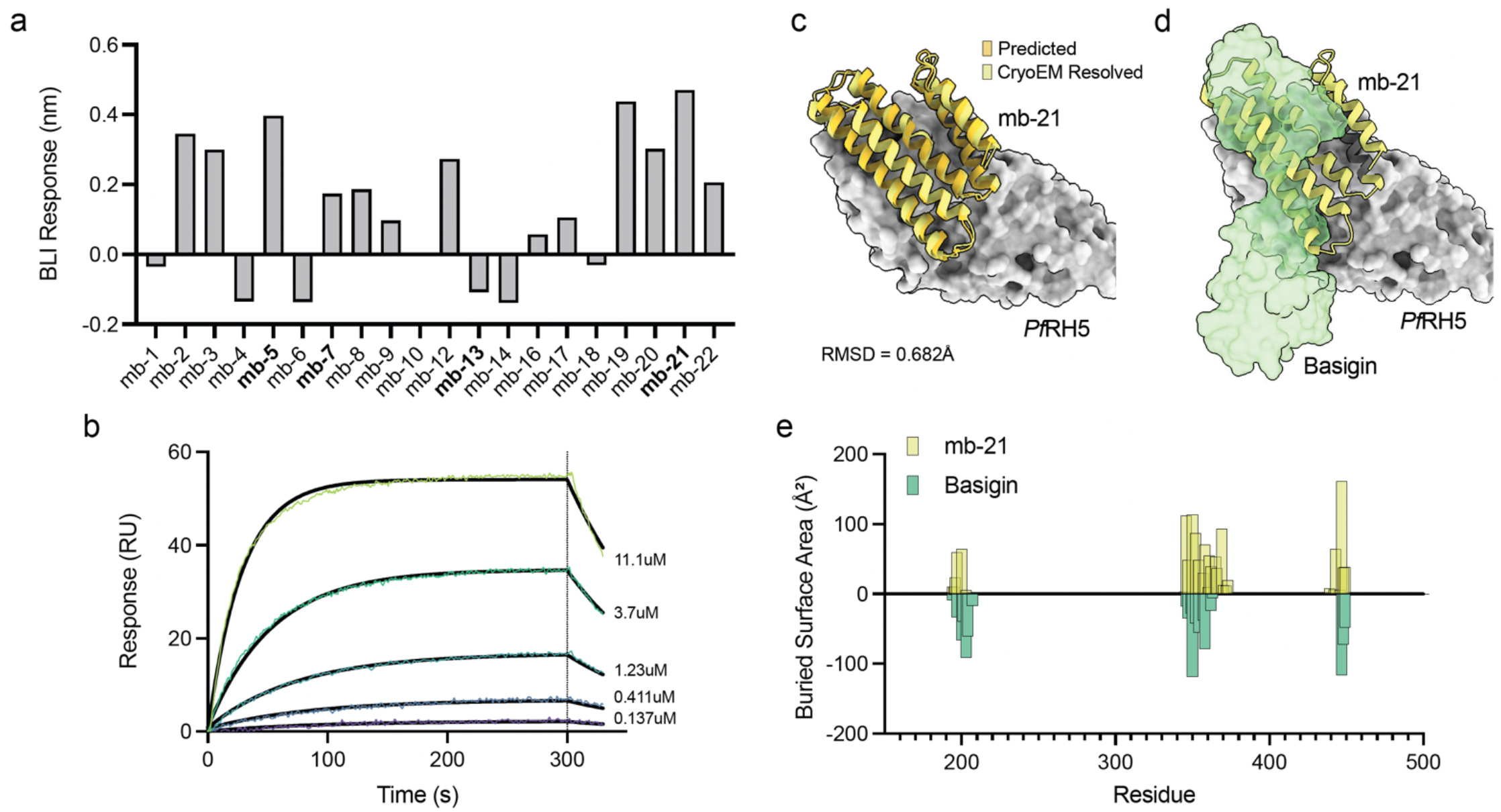
Designed minibinders bind RH5 in solution at the designed basigin-interacting interface. (**a**) BLI response amplitude for each purified design against recombinant RH5 (*n* = 1). Negative values arise from reference subtraction and indicate no binding. (**b**) SPR sensorgrams for the binding designs (*n* = 1). (**c**) Experimental CryoEM structure of mb-21 (yellow) complexed with *Pf*RH5 (grey). Predicted mb-21 is additionally overlaid and indicated in orange. (**d**) The same mb-21 and *Pf*RH5 complex superposed with the erythrocyte receptor basigin (green surface). (**e**) Per-residue buried surface area (BSA) on *Pf*RH5 for each minibinder and for basigin.

### CryoEM localizes minibinder binding to the RH5 invasion interface

With solution binding to RH5 established, we next sought direct structural evidence that the minibinders engage the interface in the designed geometry. Cryo-electron microscopy (CryoEM) was performed on minibinders mb-5, mb-7, and mb-21 bound to RH5 (***Figure 2c, d, S5***). Single-particle reconstructions yielded overall resolutions of 3.09 Å (mb-21), 3.31 Å (mb-7), and 3.32 Å (mb-5) (***Figure S4, Table S3***). For all three leads, the experimental structure superposed on the computational design model with sub-Ångström backbone agreement (Cα RMSD 0.56–0.72 Å), confirming that each minibinder adopts its intended fold and docking geometry (Figure S5, Table S5). RH5 was uniformly well-resolved across all three complexes; local resolution at the minibinder interface varied (best for mb-21, weakest for mb-7). Buried interface residues, buried surface area (BSA) (***Table S4***), and overall minibinder architecture were unambiguously interpretable, though precise rotamer-level contacts were not. All three independently designed minibinders bind on regions of RH5 that the receptor basigin contacts (amino acids ∼195–205, ∼345–370, and ∼440– 450) (***Figure 2d, e, S5***).

This demonstrates that BindCraft constrained against the three RH5 hotspot residues (449, 358, 201) reproducibly recovers binders that occlude the basigin surface across diverse scaffolds (Figure S6). Together, our kinetic and binding data establish that seven designs engage RH5 directly and CryoEM demonstrates mb-5, mb-7 and mb-21 bind to RH5 at the expected, designed basigin interface.

### Lead minibinders exhibit nanomolar potency in blood-stage screening assays

Having established that the minibinders are expressed, bind RH5, and structurally occlude the basigin site as designed, we asked whether this blockade translates into potent inhibition of parasite growth *in vitro*. To assess the inhibitory activity of the minibinders, we used a 72-hour asexual blood stage phenotypic growth assay that has been used in the discovery of many small molecule drug candidates over the past two decades (2). The 72-hour assay produces better signal to noise than a shorter, synchronized growth inhibition assay (GIA) generally utilized by the immunology field, and has been more efficient for screening thousands of candidates. Compounds discovered with this assay and which have progressed to human trials where they have shown relevant antimalarial efficacy have included KAE609 (Cipargamin) and KAF156 (Ganaplacide) (2, 26). The assay is thus able to be highly predictive of therapeutic efficacy. Here, we used luciferase-expressing *P. falciparum Dd2* (*Dd2*-Luc) instead of the traditional SYBR green method (27) and tested the minibinders in 11-point dose-response format. Dose-response measurements across 19 minibinders identified seven with sub-micromolar potency (***Figure 3a***). For subsequent analyses, we classify a minibinder “potent” via the 1µM threshold as it is a common threshold used in high-throughput drug screening programs (***Table S6, Figure 3a***). The most potent three minibinders (mb-5, mb-21, mb-7, bolded) had mean IC_50_ values below 201nM, with one of the candidates, mb-5, reaching IC_50_ of 107 ± 13nM. Compared in parallel in the same assay, these potencies were competitive to a known anti-RH5 monoclonal antibody, R5.004 (43 ± 6nM). Four additional candidates showed nanomolar activity below 309nM (mb-2, mb-12, mb-19, mb-8), while three had low micromolar activity values (mb-1, mb-17, mb-20). The remaining minibinders displayed no measurable inhibition at the highest concentrations tested (Supplementary Table S6).

**Figure 3.**
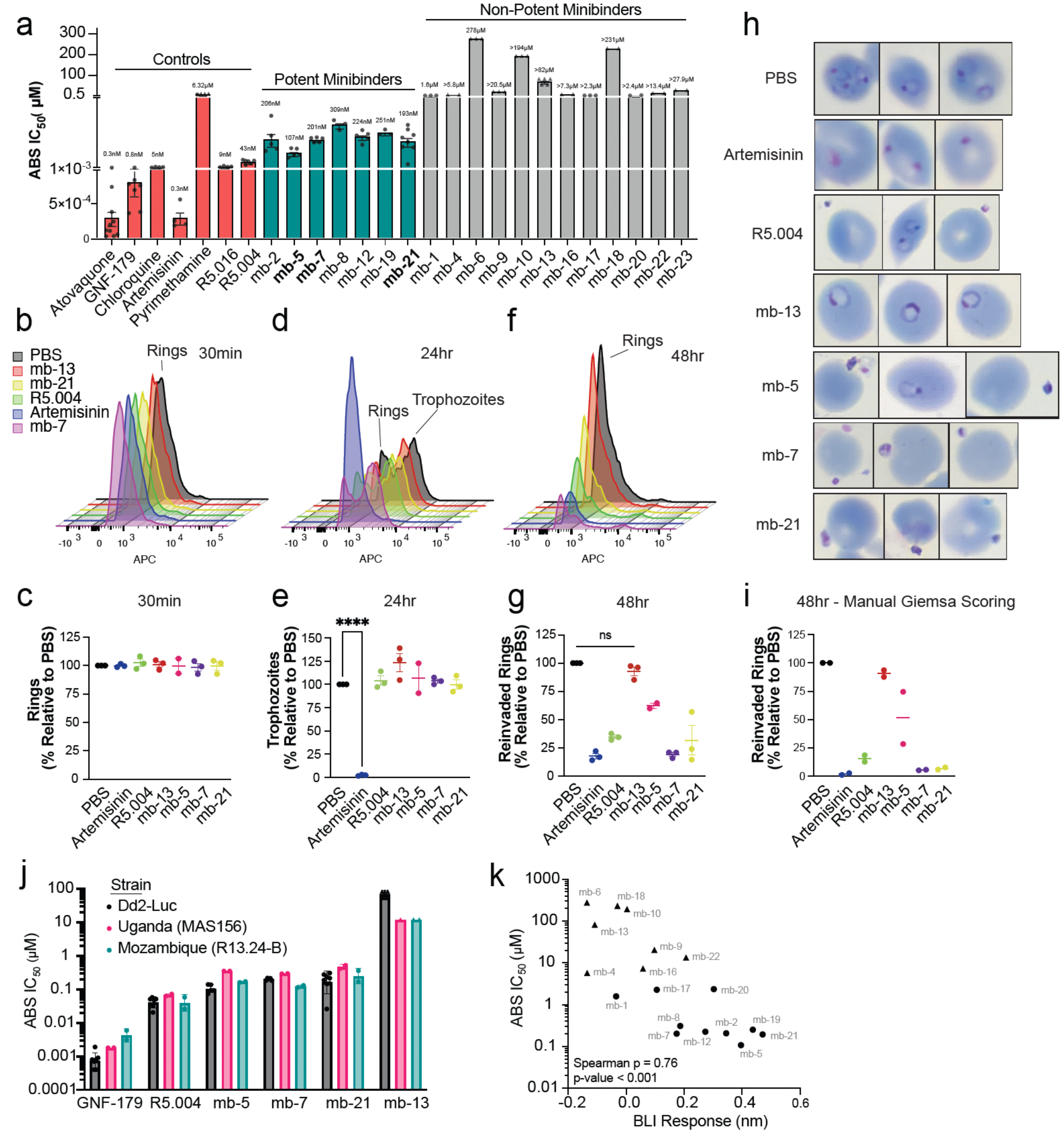
*De novo* minibinders inhibit blood-stage *Plasmodium falciparum* with potency comparable to benchmark anti-RH5 monoclonal antibody R5.004. (**a**) ABS IC_50_ values for 19 designed minibinders in *P. falciparum Dd2* luciferase growth-inhibition assay (72 h) with small-molecule antimalarial controls and anti-RH5 monoclonal antibody controls R5.004, R5.016 (potent binders/molecules *n* ≥ 3; non-potent binders *n* ≥ 2). Bars = arithmetic mean, error bars = SE; triangles = IC_50_ not interpolated below maximum tested concentration. (**b, d, f**) Stage-progression by flow cytometry: double-synchronized cultures treated with minibinders mb-5, mb-7 or mb-21, R5.004, artemisinin, an inactive minibinder mb-13, or PBS were stained for DNA at 30 min, 24h, and 48h. (**c, e, g**) Quantification of % parasitemia of ring-stage parasites (30 min, 48hr) and trophozoites (24hr) relative to PBS control at a given time point via flow cytometry (*n* = 3 for all conditions except mb-13). Bars = mean, error bars = SE. Statistical comparisons performed via one-way ANOVA with Dunnett’s comparison test comparing all means to PBS control. A table of statistical comparisons at all time points is summarized in ***Table S10***; full flow cytometry gating strategy in ***Figure S9***. (**h**) Representative Giemsa-stained thin smears at 48 hrs post-synchronization. (**i**) Manual microscopy quantification of % parasitemia of reinvaded ring-stage parasites at 48hr relative to PBS control via Giemsa-stained slides (*n* = 2). Bars = arithmetic mean, error bars = SE. (**j**) ABS IC_50_ for Dd2-Luc compared to field isolates from Uganda (MAS156) and Mozambique (R13.24-B) for a subset of minibinders (*n* = 2). Bars = mean, error bars = s.d.; triangle points = IC_50_ greater than highest concentration tested. (**k**) ABS IC_50_ versus BLI response for all screened minibinders. Higher BLI response correlates with lower IC_50_ (Spearman ρ = -0.76, p < 0.001, n = 18). Error bars = SE; triangles = IC_50_ greater than highest concentration tested.

To ask whether potency could be rationalized from binder and interface features, we compared sequence, structural, and simulated interface-dynamic properties of potent and non-potent minibinders. The clearest difference between the groups was binder length, with all seven potent designs falling within a narrow 135–143 residue window, whereas 11 of 12 non-potent designs are shorter (77–125 residues) (***Figure S7, Table S7***). However, no feature survived multiple-testing correction, including residue-level contact occupancy across the RH5 surface (***Figure S6***). Establishing structure–activity relationships for this class will likely require larger panels.

### Designed minibinders are potent against recent, field-relevant *P. falciparum* isolates bearing common *Pf*RH5 mutations

We next sought to validate that our minibinders retain activity against *P. falciparum* strains currently circulating in the field. Two recent clinical isolates, R13.24-B (Mozambique, 2024) and MAS156 (Uganda, 2018) were initially cultured in vitro with human serum and then gradually transitioned to AlbuMAX II supplementation. Whole genome sequencing of these culture-adapted field strains confirmed that R13.24-B bears three nonsynonymous single nucleotide variants (Y147H, H148D, C203Y) in RH5 relative to the *Pf*Dd2 reference genome, while MAS156 only has C203Y. These RH5 haplotypes are common in natural parasite populations, with C203Y representing approximately 86% of all 23,035 QC-passing samples in the Pf8 dataset of worldwide *P. falciparum* genomes (28) (***Methods***), while Y147H + H148D + C203Y represents ∼4% and occurs most frequently in Eastern Africa. Importantly, the *Dd2*-Luc screening strain has RH5 haplotype lacks the circulating C203Y mutation which lies adjacent to a targeted hotspot, residue 201 (***Table S8***). We took a subset (nine) of the minibinders tested in *Dd2*-Luc to test against these field isolates. Our results demonstrate that every potent minibinder tested against *Dd2*-Luc retained nanomolar inhibition against MAS156 and R13.24-B, while every non-potent minibinder remained inactive (***Figure 3j, Table S9***). Our monoclonal antibody control, R5.004, also retained similar efficacy against MAS156 and R13.24-B (68 ± 5 nM and 42 ± 16 nM, respectively), in line with prior data where R5.004 was similarly potent across seven laboratory and field strains bearing a variety of RH5 haplotypes (5). This suggests that RH5 mutations present in field isolates do not abrogate minibinder potency.

### BLI screening and SPR kinetics predict *in vitro* blood-stage phenotypic potency

To ask whether the primary BLI binding screening predicts blood-stage activity, we compared the equilibrium response (nm) of each design with its blood-stage IC_50_ against Dd2-Luc (Figure 3k). Across the 18 minibinders for which both measurements were available, BLI response magnitude was strongly and significantly correlated with potency (Spearman ρ = -0.76, p < 0.001). Designs producing a substantial RH5 binding response were uniformly potent (IC_50_ < 1µM), whereas those with responses at or below baseline were inactive (IC_50_ > 1µM). The correlation is driven predominantly by this separation between binding and non-binding designs rather than by graded differences among binders. Six of the seven <1µM designs bound RH5 by BLI, SPR or both, while eleven of the twelve designs with >1µM yielded no detectable binding signal in BLI and SPR (***Figures 2a, 2b, S3***). Within the potent set, IC_50_ values clustered near ∼0.2 µM and were not further resolved by BLI response magnitude nor kinetic measurements. Notably, mb-7 (one of the three designs resolved by CryoEM) produced only a low-amplitude BLI response yet bound detectably by SPR, potentially indicating that rapid association and dissociation can suppress apparent BLI signal without indicating an absence of binding. Additionally, mb-20 bound RH5 in BLI and SPR but was not potent while mb-8 was potent despite only modest BLI response and lack of reliable SPR kinetic measurements. We additionally validated that potent minibinders, when biotinylated for SPR, remain potent in blood-stage phenotypic assays while non-potent minibinders remain non-potent (***Figure S8***). BLI response magnitude therefore provides a reliable, high-throughput predictor of whether a design will inhibit blood-stage replication, though not how quantitatively potent it will be.

### Minibinders block merozoite to ring transition, not intracellular replication

Because these minibinders are designed to occlude the RH5-basigin interface required for merezoite invasion, we expect that they would block parasite replication at the schizont-to-ring transition (***Figure 1a, b***). To confirm the stage of inhibition, we double-synchronized cultures of *Dd2*-Luc with 5% D-sorbitol to obtain a highly synchronous ring-stage population, as previously described (29). Parasites were then co-incubated with ≈10 x IC_50_ of mb-5, mb-7, mb-21, or mb-13 minibinder (non-potent minibinder, negative control), PBS (negative control), artemisinin (positive control), or R5.004 (reference mAb, positive control). As parasites progress from rings to trophozoites (24h) and trophozoites to schizonts (a further 24h), DNA content increases geometrically before release of up to 32 merezoites per schizont for reinvasion ≈48h post-synchronization. Because erythrocytes lack DNA, we tracked parasite life cycle progression by live-cell flow cytometry using SYTO61 DNA staining (***Figure 3b, c***).

At 30 minutes post-incubation, all cultures were highly synchronized at ring-stage, marked by a single, low-APC peak (***Figure 3b***). At 30 minutes, there were no significantly different proportion of ring-stage parasites relative to PBS control (***Figure 3c***). At 24 hours, R5.004- and minibinder-treated parasites progressed normally to trophozoites, indicated by a broad, secondary peak with higher APC signal non-significantly different from PBS control (***Figure 3d***). This contrasted with artemisinin, an all-stage antimalarial, that sharply blocked the ring-to-trophozoite transition (*p* <0.0001) (***Figure 3e***). By 48 hours, when schizonts had ruptured and merezoites re-invaded to form new rings, all treatments except for non-potent mb-13 demonstrated a significantly reduced ring population (***Figure 3f, g***). This merezoite-to-ring transition blockade via R5.004, mb-5, mb-7, and mb-21 is consistent with the intended mechanism of inhibiting RH5-basigin interaction necessary for merezoite reinvasion. A summary of statistical comparisons is available in ***Table S10***.

Manual counting of Giemsa-stained smears at 48h supported the observations from our flow cytometry data (***Figure 3h, i, Table S10***): mb-5, mb-7, and mb-21-treated cultures contained extracellular puncta consistent with non-viable merozoites unable to re-enter erythrocytes, mirroring the phenotype of RH5-targeting mAb R5.004. Conversely, mb-13-treated cultures displayed a healthy ring-stage population indistinguishable from PBS-treated control. This stage-specific blockade via minibinders is consistent with our binding and kinetic results, further corroborating the intended mechanism of RH5-basigin inhibition.

## Discussion

Antimalarial discovery has long relied on two main paradigms. Small-molecule programs must convert a validated target into a specific chemical hit, a step that often stalls even when the biology is well understood and that demands extensive biochemical and genetic validation to confirm on-target action. Blood-stage biologics have depended on vaccines and reverse vaccinology, eliciting growth-inhibitory antibodies against RH5 through immunization. Both are effective but inherently slow. Here, we demonstrate a third route: target-based *de novo* protein design, in which the interface to be blocked is specified *a priori* and inhibitors are generated directly against it, without immunization, library selection, or a starting scaffold.

We generated a single first-generation panel of approximately 1500 computationally designed proteins, of which 19 were chosen to be expressed and purified for binding characterization. Subsequent screening of all 19 designs revealed 7 potent inhibitors of *P. falciparum* replication, with confirmed merozoite reinvasion blockade at the intended epitope with sub-Angstrom agreement to what was computationally predicted. Critically, while such hit rate (7/1500) is analogous to the commonly seen hit rates of <1% in compound library screening programs, the fact that screening can be performed largely *in silico* greatly streamlines potent lead identification. In theory, potent binders can also be discovered even against interfaces that are only partially characterized, reflecting the maturation of BindCraft’s computational pipeline and high-throughput screening capabilities. Aside from the modest inhibition recently reported for *de novo* binders against the AMA1-RON moving-junction complex (30), this is, to our knowledge, the first demonstration of potent de novo-designed inhibition of malaria parasite growth.

Our lead minibinders (mb-5, mb-7, and mb-21; approximately 100-200 nM IC_50_) are competitive in potency to the benchmark antibody R5.004 assayed here and even outperform recently reported anti-RH5 nanobodies (8) (lowest IC_50_ ≈ 109 nM), a level normally reached only through immunization and selection. Because both R5.004 and R5.016 appeared somewhat more potent in our assay than in published growth-inhibition assays (5, 6), potentially owing to asynchronous parasites and longer incubation of our assays, we anchored cross-assay comparisons to the shared R5.004 control. On this basis, our lead was more potent than roughly two-thirds of a 247-member panel of vaccine-elicited human anti-RH5 monoclonal antibodies (6), despite requiring no affinity maturation. Our limited panel of minibinders do remain one to two orders of magnitude less potent than historically important small molecules including artemisinin and atovaquone. This is consistent with the fact that the K_D_ of our most potent minibinders to RH5 is low-micromolar compared to nanomolar of matured human anti-RH5 monoclonals. We regard these gaps as the expected starting point for an unmatured design campaign rather than a ceiling, and as a benchmark of the necessary improvements to antiparasitic and binding kinetics that remain. Importantly, the minibinders act through a mechanism orthogonal to every deployed antimalarial. By occluding the RH5-basigin interface, they arrest invasion such that their activity is unaffected by the resistance mutations that erode current frontline antimalarials, including pyrimethamine, artemisinin, and atovaquone. Because RH5 is under strong functional constraint and among the most conserved blood-stage antigens, our potent designs expectedly retained activity against circulating clinical isolates from Uganda (MAS156) and Mozambique (R13.24-B).

Beyond RH5, the same pipeline can be directed at other validated malaria targets, including CyRPA and the broader PCRCR complex, the AMA1-RON2 junction, and even pre-erythrocytic ligands such as CSP, enabling combination and multi-stage strategies and offering a rapid, generalizable route to targets that have resisted conventional drug and vaccine development. Realizing this will firstly require improvements in potency and binding affinity, with off rates particularly the clearest targets for improvement. We additionally hypothesize that conformational rigidity of the minibinder itself can be scrutinized further in future designs. Affinity maturation, multivalent binders with linkers, half-life extension, and larger screening panels, all more tractable on these *de novo* designed scaffolds than on antibodies, offer well-precedented routes to improving potency and developability. Additionally, *in vivo* efficacy in humanized-mouse and non-human-primate models and careful assessment of the immunogenicity of these non-natural sequences, need to be better understood. These caveats and limitations of our current study notwithstanding, our results establish that the essential RH5-basigin interaction can be blocked by molecules that are designed rather than discovered and chart a plausible path for *de novo* protein generation toward a new class of antimalarial biologics.

## Materials and Methods

### Computational design and validation

*De novo* minibinders were generated against PfRH5 (PDB 4WAT) using BindCraft (12)with hotspot residues 449, 358, and 201, identified from interface analysis of the RH5–basigin complex (PDB 4U0Q). Designs were filtered to length 75–175 residues, helicity-favoured, cysteine-free, with i-PAE < 0.35, pLDDT > 0.86, Rosetta ΔG < −45 kcal mol^−1^, and ΔG/ΔSASA < −2.0 kcal mol^−1^ Å^−2^. Top candidates were refined with ProteinMPNN [Dauparas, J. et al. 2022] and independently validated by AlphaFold3 (14) (two model seeds per design) and Boltz-2(17) (25 recycling steps, one diffusion sample). Designs with ipTM < 0.7 or detected steric clashes were excluded. Full pipeline configuration, JSON templates, and SLURM scripts are deposited at github.com/jinichlab/RH5-minibinders; Supplementary Methods §1 lists exact parameter sets.

### Minibinder expression and purification

Twenty-four design genes were codon-optimised for *Escherichia coli* (Integrated DNA Technologies) and cloned by Golden Gate assembly into the LM670 expression vector (gift of T. Bethel, UCSD; original source D. Baker, UW), encoding a C-terminal hexahistidine tag. Constructs were transformed into chemically competent BL21 cells in 96-well format and expressed by 20 h autoinduction at 37 °C. Cells were lysed with B-PER reagent (Thermo Scientific) supplemented with DNase I and PMSF, and proteins were purified by Ni-NTA magnetic-bead capture (PureProteome, Millipore) followed by imidazole elution. Lead candidates were re-expressed at 100 mL scale (LB, IPTG induction at 18 °C overnight), purified on Ni-NTA resin (HisPur, Thermo Scientific), and polished by size-exclusion chromatography on a Superdex 200 Increase 10/300 GL column (Cytiva). Monomeric peak fractions were pooled and quantified by DC Protein Assay (Bio-Rad). Detailed buffer compositions, plate formats, and scale-up protocols are in Supplementary Methods.

### R5.004 construct preparation, expression, and purification

The R5.004 heavy chain (HC) and light chain (LC) variable domain sequences of the potency antibodies were obtained from the RCSB Protein Data Bank. Gene fragments encoding the HC and LC variable regions were synthesized by Integrated DNA Technologies (IDT, San Diego, CA, USA) and cloned into pAbVec mammalian antibody expression vectors containing the human IgG1 heavy chain constant region or the human kappa light chain constant region using Gibson assembly. All expression constructs were verified by whole-plasmid sequencing (Quintara Biosciences, San Diego, CA, USA). Validated HC and LC plasmids were transiently expressed in Expi293F cells (Thermo Fisher Scientific). Cells were maintained in Expi293 Expression Medium at 37°C in a humidified incubator with 8% CO_2_ and orbital shaking at 125 rpm and routinely cultured at densities ranging from 3 × 10^5^ to 5 × 10^6^ cells/mL. For transient transfection, HC and LC plasmids were mixed at a 2:1 (w/w) DNA ratio and complexed with FectoPRO Transfection Reagent (Sartorius) in Opti-MEM Reduced Serum Medium (Thermo Fisher Scientific) according to the manufacturer’s recommendations. Transfections were performed at a cell density of 2.5–3.0 × 10^6^ cells/mL. Twenty-four hours after transfection, cultures were supplemented with 3 mM valproic acid (Sigma-Aldrich) and 4 g/L D-glucose. Culture supernatants were harvested 5 days post-transfection, clarified by centrifugation at 4,000 × g for 30 min, and filtered through a 0.22-μm membrane filter. Clarified supernatants were incubated with Protein G resin (Cytiva) overnight at 4 °C with gentle agitation. After washing, bound antibodies were eluted using IgG Elution Buffer (Thermo Fisher Scientific) and immediately neutralized with 1 M Tris-HCl (pH 8.0). The eluates were concentrated and buffer-exchanged into phosphate-buffered saline (PBS) using Amicon Ultra centrifugal filter units (Millipore), aliquoted, and stored at −20 °C until further use.

### Blood-stage phenotypic screening assay with *P. falciparum Dd2-Luc*

A luciferase-expressing *Dd2* strain(27) - was maintained at 37°C in RPMI 1640 supplemented with 1µM blasticidin S HCl, 0.5mM hypoxanthine, 42µM gentamicin, and 0.25% (w/v) AlbuMAX II (GIBCO), at 2% hematocrit in O+ human erythrocytes (Scripps Research Institute blood bank) under low-oxygen atmosphere (1% O_2_, 5% CO_2_, 94% N_2_). Asynchronous cultures were seeded at 0.5% parasitemia and 0.5% hematocrit in 384-well plates and co-incubated for 72 h with serial dilutions of minibinders or control molecules prepared by automated pipetting (VOYAGER/Assist Plus, Integra Biosciences). Parasite viability was measured by luminescence after addition of BrightGlo reagent (Promega) on a PHERAstar FSX reader (BMG Labtech). Each condition was performed as technical quadruplicates; the assay was repeated with three or more biological replicates for minibinders with defined IC_50_ values, and with at least two biological replicates for inactive minibinders with IC_50_ values that could not be interpolated below the maximum tested concentration. IC_50_ values were determined using CDD Vault’s built-in four-parameter logistic curve fitting (unconstrained), described below; 0% response was defined by infected, Atovaquone- (Sigma Aldrich) and GNF179-treated (MedChemExpress) wells at 0.1µM. Conversely, 100% response was defined by infected, PBS-only wells. R5.004 and R5.016 were included as known, reference anti-RH5 monoclonal antibodies.

### Blood-stage phenotypic screening assay with *P. falciparum* field isolates

Field isolates from Uganda (MAS156) and Mozambique (R13-24.B) were maintained at 37°C in RPMI 1640 supplemented with 1µM blasticidin S HCl, 0.5mM hypoxanthine, 42µM gentamicin, and 0.25% (w/v) AlbuMAX II (GIBCO), at 4% hematocrit in O+ human erythrocytes (Scripps Research Institute blood bank) under low-oxygen atmosphere (1% O_2_, 5% CO_2_, 94% N_2_) on an orbital shaker to facilitate growth. Asynchronous cultures were seeded at 1% parasitemia and 2% hematocrit in 96-well V-bottom plates and co-incubated for 96h on an orbital shaker with serial dilutions of minibinders or control molecules prepared by automated pipetting (VOYAGER/Assist Plus, Integra Biosciences). Parasite stage and viability and was measured by flow cytometry (BD FACS Canto II) detecting SYTO61 (Invitrogen) signal in the APC channel as previously described (31). Infected, GNF-179 (MedChemExpress) and PBS-treated wells were included as positive and negative controls, to define as 100% inhibition and 0% inhibition, respectively. R5.004 was included as a known, reference anti-RH5 monoclonal antibody. Each condition was repeated with two biological replicates, except for mb-13 and mb-19 in Uganda (MAS156) strain. IC_50_ values were determined using CDD Vault’s built-in four-parameter logistic curve fitting (unconstrained), described below.

### Stage progression assay

Cultures were double-synchronized to ring stage by 5% sorbitol lysis and treated with minibinder, control antibody R5.004, or PBS at 10 × EC_50_, at 2% parasitemia and 2% hematocrit in 12-well format. At 0.5, 20, and 48 h post-treatment, 10µL aliquots of the culture were stained with 2µM SYTO61 (Invitrogen) and analyzed by flow cytometry on a BD FACS Canto II in the APC channel. Infected erythrocytes were identified by sequential gating on size/granularity, singlets, and APC positivity above an uninfected-control threshold; additional stage identification, alongside a summarized gating strategy are shown in Supplementary Figure S9. Each condition was repeated a total of three biological replicates, except for two biological replicates for mb-5.

### Giemsa-Stained Microscopy

Parasitemia and developmental morphology (rings, trophozoites, schizonts, gametocytes, extracellular merozoites, dead puncta) were manually scored on Giemsa-stained thin blood smears at 48 hours post-treatment by a blinded observer, counting ∼410-770 total red blood cells per sample across two biological replicates. The raw count reads are provided in Supplementary Table S10.

### Statistical analysis

IC_50_ values were determined using CDD Vault’s built-in four-parameter logistic curve fitting (unconstrained). Minibinders with IC_50_ > 1µM were assayed in a minimum of two independent biological replicates, and those with IC_50_ < 1µM in three or more, reflecting a threshold that clearly classified potent binders from non-potent binders as assessed by potency and binding data. Additional replicates were not pursued for non-potent minibinders as no downstream claims depended on their precise potency. Each independent assay comprised two to four technical replicates. Reported IC_50_ values represent the arithmetic mean of per-biological replicate IC_50_ values, with standard error of the mean calculated across biological replicates.

All additional statistical analyses and data visualization were performed using GraphPad Prism (v11.0.0) or Python.

### Protein Expression and Purification for Cryo-EM, BLI and SPR

RH5.1 and RIPR proteins were a gift from Simon J. Draper, University of Oxford, UK(32). The wild-type 3D7 strain sequence of CyRPA was mutated to remove N-glycan sequons by mutating S/T to A and subcloned into a pcDNA3.1 vector with a c-terminal Twin-strep purification tag by Genscript and an n-terminal 18-amino acid signal peptide derived from human serum albumin, USA and expressed using the Expi293 expression system (Thermo Fisher Scientific) according to the manufacturer’s instructions. CyRPA protein was affinity purified using Strep-Tactin®XT 4Flow® resin (IBA Lifesciences) and further purified by size exclusion chromatography into TBS. For the production of RH5 for BLI, the codon optimized RH5.1 sequence was cloned into a pcDNA3.1 vector with a c-terminal EPEA Ctag and an n-terminal 18-amino acid signal peptide derived from human serum albumin. This construct was expressed using the Expi293 Pro expression system (Thermo Fisher Scientific) according to manufacturer instructions. RH5 protein was purified from supernatant using CaptureSelect− C-tagXL Affinity Matrix (Thermo Fisher Scientific) using 4mM SEPEA peptide for elution. The protein was further purified using size exclusion chromatography into TBS pH 7.14. Purified protein was snap frozen and stored for future use. For the production of R5.011 Fab for Cryo-EM and BLI, the published heavy and light chain Fv sequences were synthesized and subcloned into Abvec IgG1 vectors by Genscript, USA(5). Heavy and light chains were co-expressed in a 1:1 ratio using the Expi293 expression system (Thermo Fisher Scientific) according to the manufacturer’s instructions. R5.011 Fab was then purified using CaptureSelect− CH1-XL Affinity Matrix (ThermoFisher Scientific) according to manufacturer’s guidelines. The resultant fab was then buffer exchanged into TBS pH 7.14 and frozen for future use.

### Cryo-EM sample preparation and data collection

Individual RCR-complex components were mixed at a 1:1 molar ratio in TBS and incubated for 10 min at RT. Minibinders were added to the RCR-complex mixture at 10-fold molar excess of minibinder at a final total protein concentration of 0.8 mg/mL in TBS and incubated for 10 min at RT. For the mb-7 and mb-21 minibinder complexes, R5.011 Fab was also added at a 1:1 molar ratio of fab to RCR complex to act as a fiducial for particle orientation sampling. 3 μL of each complex was added to UltrAuFoil R 1.2/1.3 300 mesh gold grids which had been plasma washed by glow discharging with Leica Coater ACE200 for 30 s at 10 mA. The sample was then plunge frozen in liquid ethane using Vitrobot Mark IV (Thermo Fisher) after incubation for 5s, blotted for 4 s at 4 °C, 100% humidity. The grids were then stored in liquid nitrogen for data collection.

Grids were loaded into a Glacios 2 microscope (Thermo Fisher Scientific) operating at 200 kV with a Falcon 4i direct electron detector, Automated data collection for all complexes was carried out using EPU (Thermo Fisher Scientific) at a nominal magnification of 190,000, with a total exposure dose of ∼45 e-/Å2 at a pixel size of 0.718 Å with a nominal defocus range of -0.8 to -2 μm. Full data collection parameters for each map and model are shown in Supplementary Table S3.

### Cryo-EM data processing and model building

All datasets were processed using cryoSPARC(33). Dose-weighted movie frame alignment was carried out using Patch motion correction in cryoSPARC live to account for stage drift and beam-induced motion. The contrast transfer function (CTF) was estimated using Patch CTF in cryoSPARC live. Micrographs with a CTF fit >10 Å were excluded. Particles were selected from micrographs using blob picker and extracted at a pixel size of 2.872 Å. 2D classification was performed and particles from good classes were used for ab-initio reconstruction, followed by heterogenous refinement. The particles from the best class from heterogenous refinement were re-extracted at full pixel size (0.718 Å) and subjected to 3D classification with 10 classes and a filter of 4Å. Particle were selected from 3D classes with good resolution, minimal orientation bias and high minibinder occupancy. The resultant particle stack and the best map were used for a non-uniform refinement job (34), followed by global CTF correction and local CTF correction, and subjected to a final local refinement job using a mask around RH5 and the minibinder (generated in ChimeraX (35)) to yield the final map for deposition and final refinements. For initial model building and figure making, the map was postprocessed by EMready2 (36).

The previously predicted models of minibinders, an AlphaFold3 predicted model of R5.011 fab and chains B (CyRPA) and C (RH5) from PDB ID:8CDD (19) were fitted into the EMready2 map in UCSF ChimeraX. A model for RIPR was not built as this region was not included in the local refinement mask. The models were manually adjusted and refined using Coot (37) and further refined through real-space refinement in Phenix (38)using the final local-refinement map output. Buried surface area (BSA), and hydrogen bonds were analyzed using the PDBePISA server (39). Structural figures were generated using UCSF ChimeraX(35).

### Bio-Layer Interferometry

Minibinders were screened for RH5 binding on an Octet® RED96e system (Sartorius). Octet® FAB2G Biosensors (Sartorius) were first dipped into Kinetics buffer (PBS + 0.01% BSA + 0.002% Tween 20 pH 7.14). Sensors were then dipped into wells containing R5.011 Fab at 10 μg/mL in Kinetics buffer for 2 minutes, followed by RH5 protein at 60 μg/mL in Kinetics buffer for 2 minutes and then wells containing each minibinder at 10 μg/mL in Kinetics Buffer for 2 minutes. Between each step, sensors were dipped into Kinetics buffer to establish a baseline. Data was processed in the Octet® Analysis Studio software (Sartorius). Minibinder sensograms were referenced using a sensor that was not loaded with RH5. The response was recorded at the end of the minibinder association phase and figures were made using GraphPad Prism 10.6.1.

### Surface Plasmon Resonance

A subset of monobiotinylated avi-tagged minibinders were selected for SPR experiments using the Carterra LSA^XT^ (Carterra). A Carterra SAP biosensor chip (#4295) was activated with several buffer injections of Kinetics buffer (PBS + 0.01% BSA + 0.002% Tween 20 pH 7.14). Biotinylated minibinders were printed onto one quadrant of the chip using the 96-channel print head and captured onto the streptavidin-coated surface. The surface was then stabilized using several buffer injections and then a capture kinetics protocol was performed. A 6-point three-fold dilution series of RH5.1 was prepared in kinetics buffer, starting at 33.3 μM concentration. Increasing concentrations of RH5.1 in kinetics buffer were ran over the whole chip without regeneration between steps with 5-minute association and 15-minute dissociation periods. Kinetics data was processed in the Carterra Kinetics analysis software. Data was referenced and baseline corrected and aligned to the start of each injection series. The 33.3 μM injection was excluded as sensogram traces at this higher concentration showed features consistent with secondary binding or dimerization, leaving a 5-point dilution series. Kinetic parameters were then calculated by fitting the data to a 1:1 Langmuir model. The data was also subjected to steady state analysis in the Carterra Kinetics software to generate equilibrium K_D_ values. To obtain robust, comparable constants under the 1:1 model, the analysis window was restricted to the association phase plus the first 30s of dissociation to capture the dominant initial off rate, as there was evidence of biphasic dissociation for some minibinders during the full dissociation time. Reported kinetic constants are therefore apparent parameters. All reported values are the mean ± s.d. of two technical replicates.

### Reagent and data availability

Reagents and design details are available from the corresponding author upon reasonable request.

## Data Availability

Design sequences, computational metrics, and analysis scripts will be made available upon publication. CryoEM maps and models for the mb-5, mb-7, and mb-21 structures will be uploaded onto the PDB and EMDB servers and made public upon publication. Additional experimental data including pharmacokinetic profiling is ongoing and will be included in the final dataset.

## Acknowledgments

Part of this work was supported by the HHMI Hanna Gray Fellowship (GT16787), and the NIH FIRST Faculty Program (NCI 1UA54CA272220). This research was additionally funded by the National Institutes of Health, grant numbers R01 AI169892.

Molecular graphics and analyses performed with UCSF ChimeraX, developed by the Resource for Biocomputing, Visualization, and Informatics at the University of California, San Francisco, with support from National Institutes of Health R01-GM129325 and the Office of Cyber Infrastructure and Computational Biology, National Institute of Allergy and Infectious Diseases.

