## Supplemental Information for "*De novo* minibinders targeting RH5 achieve nanomolar inhibition of blood-stage malaria replication"

***Cited Literature***........................................................................ *Error! Bookmark not defined.*

#### Supplementary Methods

##### 1. Target Selection and Hotspot Identification

The target structure for binder design was PfRH5 (PDB 4WAT, resolution 2.18 Å) (1). Interface analysis was performed on the RH5–basigin complex structure (PDB 4U0Q, resolution 3.1 Å) (2) to identify the residues that mediate basigin binding. Three critical residues were selected as design constraints (RH5 residues 449, 358, and 201), all clustered near the "kite-tip" of RH5 within the basigin-binding interface. The choice of these residues reflected two convergent lines of evidence: the structural RH5–basigin contact interface (PDB 4U0Q) and the epitopes of neutralizing anti-RH5 monoclonal antibodies (below), which independently implicate the same surface.

This hotspot selection was chosen to achieve greater epitope coverage than existing neutralizing antibodies. Whereas neutralizing monoclonal antibodies such as R5.004, 9AD4, and QA1 inhibit invasion by partially overlapping the basigin-binding site (main-text ***Figure 1c***), the designed minibinders were constrained to engage all three hotspot residues simultaneously, enabling more complete steric occlusion of the RH5–basigin interaction.

##### 2. Computational Design Pipeline

###### 2.1 BindCraft design

Within BindCraft, binder sequences are generated by backpropagation through the AlphaFold2-multimer network (via ColabDesign), then redesigned with ProteinMPNN (3), re-predicted with AlphaFold2 (4), and scored on Rosetta-based interface energetics (5). We provided the *Pf*RH5 target structure (PDB 4WAT) (1) and interface hotspot residues 449, 358, and 201, and configured the run with target lengths of 75–175 residues, a helicity bias of −0.3 (favoring helical structures), and cysteine exclusion. Designs were retained that passed the following filters: i-PAE < 0.35, pLDDT > 0.86, Rosetta $\Delta G < -45$ kcal/mol, and $\Delta G/\Delta SASA < -2.0$ kcal/mol/Å$^2$.

###### 2.2 AlphaFold3 validation

Top-ranked BindCraft candidates were independently validated with AlphaFold3 (6) on the Jinich Laboratory high-performance computing cluster. A custom Python script programmatically generated AlphaFold3-compatible JSON input files for each design. The template specified the full-length RH5 target sequence (Chain A, 469 residues) with pre-computed multiple sequence alignments (MMseqs2 against UniRef90 and environmental databases); each binder sequence was substituted into Chain B. Predictions used two model seeds per design to assess consistency. Validation metrics included the interface predicted TM-score (ipTM), interface PAE, overall ranking scores, and clash detection; designs with ipTM < 0.7 or detected steric clashes were flagged for exclusion.

##### 2.3 Boltz-2 validation

As an orthogonal check using a distinct methodology, Boltz-2 (7) predictions were run on the Jinich Laboratory GPU cluster (SLURM scheduling), using 25 recycling steps per structure and one diffusion sample per design and generating full predicted-aligned-error matrices. Outputs were parsed and integrated with the BindCraft statistics for consolidated analysis.

From the filtered set, 19 representative, cysteine-free, single-domain designs were selected for experimental characterization and carried forward to expression, purification, and blood-stage screening.

#### 3. Protein Expression and Purification

##### 3.1 Gene synthesis and molecular cloning

Genes encoding the designed RH5 minibinders were codon-optimized for *Escherichia coli* and synthesized as linear DNA fragments (eBlocks, Integrated DNA Technologies). Fragments were cloned into the LM670 expression vector (Addgene #191552; deposited by David Baker, a gift of the Bethel lab, UCSD), which carries a kanamycin-resistance marker and a ccdB selection cassette flanked by BsaI sites and appends a C-terminal hexahistidine (6×His) tag for purification(8).

##### 3.2 High-throughput expression and purification

Minibinder genes were assembled by Golden Gate reaction (5 µL; 37 °C, 20 min) and transformed into chemically competent *E. coli* BL21 in 96-well format. After heat shock and 1 h recovery in LB, 25 µL of each culture was inoculated into 1 mL autoinduction medium in 96-deep-well plates and grown for 20 h (37 °C, 400 rpm). Cells were harvested (4,000 × g, 15 min) and lysed with B-PER reagent (Thermo Scientific) supplemented with DNase I and PMSF (37 °C, 800 rpm, 30 min). Target proteins were captured from cleared lysates on PureProteome Nickel Magnetic Beads (Millipore) in 96-well plates (60 min binding, 37 °C), washed four times (25 mM Tris-HCl pH 8.0, 300 mM NaCl, 30 mM imidazole), and eluted in high-imidazole buffer (500 mM imidazole). Purity was verified by SDS-PAGE, and samples were filtered for downstream HPLC and biophysical characterization.

##### 3.3 Preparative scale-up

Promising candidates were scaled up to confirm yield and purity. Selected clones were grown in 200 mL LB (37 °C) to OD600 0.8–1.0, induced with IPTG, and expressed overnight at 18 °C. Cells were harvested (4,000 × g, 15 min), resuspended in lysis buffer, and disrupted by sonication. Clarified lysate was incubated with HisPur Ni-NTA resin (Thermo Scientific) for 1 h at 4 °C, washed with Tris-based wash buffer (30 mM imidazole), and eluted (500 mM imidazole). After SDS-PAGE, proteins were polished by size-exclusion chromatography (Superdex 200 Increase 10/300 GL, Cytiva) on an ÄKTA FPLC (Fig.S1). Monomeric peak fractions were pooled and determined by using a NanoDrop spectrophotometer and subsequently quantified via HPLC (***Figure S2***).

##### 3.4 N-Terminal Avi-Tag Conjugation and In Vitro Enzymatic Biotinylation

To facilitate site-specific functionalization and oriented immobilization for downstream biophysical assays, a biotin acceptor peptide tag (Avi-tag, GLNDIFEAQKIEWHE) was genetically fused to the N-terminus of the designed minibinders, separated by a flexible poly-glycine-serine linker (GGSGG). Following recombinant expression and initial IMAC purification, the N-terminally tagged minibinders underwent in vitro enzymatic biotinylation. The reaction was catalyzed by recombinant BirA biotin ligase (Addgene, plasmid #20857, deposited by Alice Ting), which was recombinantly expressed and purified in-house. Biotinylation reactions were carried out in the presence of 200μM biotin, 5mM ATP, and 5mM $MgCl_2$ (30°C, 40 min), allowing site-specific covalent attachment of biotin to the target lysine residue within the Avi-tag sequence. Following the enzymatic reaction, the mixture was subjected to size-exclusion chromatography (SEC) via ÄKTA (FPLC) to purify the biotinylated minibinders and completely remove unreacted free biotin, BirA enzyme, and reaction byproducts prior to functional characterization(9).

#### 4. Cryo-Electron Microscopy of the RH5-Minibinder Complexes

To obtain direct structural evidence for the designed binding mode, minibinders mb-5, mb-7, and mb-21 were each reconstituted with the RH5–CyRPA–RIPR (RCR) complex and a fiducial antibody (to break pseudo-symmetry and aid particle alignment) and analyzed by single-particle cryo-electron microscopy in collaboration with the Ward Laboratory (Scripps Research; A. Ward and J. Barrett). Grids were vitrified on glow-discharged holey-carbon supports and imaged on a Titan Krios equipped with a K3 detector. Two-dimensional classification, ab initio reconstruction, and refinement were performed in cryoSPARC. Reconstructions reached overall resolutions of 3.09 Å (mb-21), 3.31 Å (mb-7), and 3.32 Å (mb-5). For all three leads, the experimental structure superposed on the computational design model with sub-Ångström backbone agreement (Cα RMSD 0.72 Å for mb-5, 0.56 Å for mb-7, and 0.68 Å for mb-21), confirming that each minibinder adopts its intended fold and docking geometry. All three occupy the basigin footprint on RH5, providing the structural basis for blockade of basigin-dependent erythrocyte invasion.

#### 5. Data Analysis and Statistics

Dose-response curves and IC50 values were computed by four-parameter logistic regression. For consolidated analysis of design metrics versus experimental outcomes, custom Python scripts merged data from three sources: (1) BindCraft design statistics (predicted binding metrics, pLDDT, PAE, Rosetta energies, and interface properties); (2) AlphaFold3 validation data (ipTM, ranking scores, minimum PAE, and clash detection); and (3) experimental data (well positions, molecular weights, and IC50 values). Designs are referred to throughout by their systematic minibinder identifiers. Reported IC50 values summarize qualified biological replicates, converted to molar units using calculated molecular weights. Statistical comparisons between top hits and other tested minibinders used Mann–Whitney U tests given the small sample sizes, and Spearman correlation assessed the relationship between computational ranking and experimental potency. Analyses used pandas, numpy, scipy, and matplotlib.

#### 6. Computational Structural Analysis of the Design Panel

##### 6.1 Potency grouping

For the subset of minibinders with complete structural models and qualified Dd2-Luc potency measurements (n = 19), designs were grouped as potent (n = 7; IC50 < 1 μM) or non-potent (n = 12) for computational comparison.

##### 6.2 Structural registration and residue-contact mapping

Predicted RH5-minibinder complexes were registered to a common RH5 reference frame using chain A of PDB 4WAT. Cα atoms matched through the biological RH5 residue map were superposed with Bio.PDB; target-registration Cα RMSD across the 19 predicted complexes ranged from 0.68 to 0.90 Å, suggesting similar folds across all binder co-folds. Binder-RH5 contacts were recomputed directly from coordinates using a 5.0 Å heavy-atom distance cutoff. For each design, the analysis recorded the RH5 footprint, binder-interface residues, residue-contact pairs, minimum inter-residue distances, footprint radius of gyration, binder standoff, and the number of design hotspots contacted. Residue-level contact fingerprints for the full design panel are shown in Figure S6.

##### 6.3 Reference-epitope comparison and interface geometry

Reference complexes were placed in the same RH5 frame and analyzed with the same 5.0 Å contact definition: basigin (PDB 4U0Q (2)), neutralizing antibodies R5.004 and R5.016 (PDB 6RCU (10)), R5.034 (PDB 8QKS (11)), and non-neutralizing R5.015 (PDB 7PHU (12)). Footprint overlap was quantified by Jaccard similarity. Basigin coverage was defined as the fraction of basigin-contacting RH5 positions recovered by a minibinder footprint. Solvent-accessible surface area was calculated with the Shrake-Rupley algorithm; total buried area was defined as the summed loss of solvent-accessible surface from RH5 and the minibinder upon complex formation. These 5 Å/Shrake-Rupley values are used for the supplementary structural comparisons below. The resulting geometry comparisons are reported in Table S7.

##### 6.4 Residue-level and scaffold-aware statistical analysis

For each of the 45 unique RH5 residues contacted by at least one minibinder in the 19-design panel, contact occupancy in potent versus non-potent designs was compared using Fisher's exact test, with Benjamini–Hochberg correction across the 45 residue-wise tests. Continuous geometry features were summarized by group medians and Cliff's delta. Group differences were evaluated by exact enumeration of the median-difference statistic at the design level and, where appropriate, after collapsing repeated designs to scaffold-level means. Benjamini-Hochberg correction was applied across the tested geometry features. Corrected significance values are reported alongside each comparison in Figure S6 and Table S7.

Scaffold-level collapsing controls for non-independence among designs sharing a backbone but does not remove an association between binder size and potency group. Binder length was examined as a potential confounder of the feature comparison. All seven potent designs fall within a narrow 135–143 residue window, whereas eleven of twelve non-potent designs are shorter (77–125 residues). Binder length alone separates the groups with Cliff's $\delta = 0.83$ (exact $p = 1.6 \times 10^{-3}$), but does not survive multiple-testing correction across the features examined. Features scaling with binder size may therefore separate the groups for reasons unrelated to potency.

##### 6.5 Model-to-experiment comparison for cryo-EM leads

For mb-5, mb-7, and mb-21, the experimental cryo-EM complexes and the corresponding computational models were independently registered through RH5 to the common 4WAT frame. Agreement was quantified using experimental-versus-predicted RH5-footprint Jaccard similarity and the angular difference between predicted and experimental coarse approach vectors. These measurements are used only as retrospective pose-validation metrics for the three experimentally solved leads and are not extrapolated to the unsolved designs. Model-to-experiment comparisons are reported in Figure S6 and Table S5.

#### 7. Molecular Dynamics Simulations of Predicted Complexes

##### 7.1 System preparation

AlphaFold-predicted RH5 minibinder complexes were prepared with PDBFixer (https://github.com/openmm/pdbfixer). Non-standard residues were replaced with standard equivalents, missing residues and heavy atoms were reconstructed, and hydrogens were added assuming pH 7.4 to match the conditions of the biochemical assays. Heterogens were retained and crystallographic waters removed. Structures were relaxed by L-BFGS minimization with hydrogen-bond constraints (convergence tolerance 100 kJ $mol^{-1}$ $nm^{-1}$; maximum 10,000 iterations), then solvated in periodic truncated boxes with a minimum solute–box distance of 15 Å using the TIP3P water model (13) and neutralized with $Na^+$ and $Cl^-$ to a physiological ionic strength of 0.15 M. Proteins were described with the AMBER ff14SB force field (14). Hydrogen mass repartitioning to 4.0 amu (15) was applied to permit a 4 fs integration timestep under pressure coupling.

##### 7.2 Molecular dynamics

Simulations were performed with OpenMM 8.2 (16) on NVIDIA GPUs (CUDA platform, mixed precision). Long-range electrostatics used Particle Mesh Ewald with a 1.2 nm real-space cutoff, and all bonds involving hydrogen were constrained. With backbone atoms (N, Cα, C) harmonically restrained at 100 kJ $mol^{-1}$ $nm^{-2}$, systems were heated stepwise to 300 K and equilibrated for 2 ns in the NPT ensemble. Restraints were then released and each system propagated for 100 ns of production under identical NPT conditions using a LangevinMiddle integrator (300 K, collision frequency 1 $ps^{-1}$), a Monte Carlo barostat (1 bar, updated every 25 steps), and a 4 fs timestep. Three independent production replicates were run per system, differing only in the random seed used to initialize the thermostat and atomic velocities. Equilibration was verified before production by monitoring potential energy, total energy, temperature, density, and box volume.

##### 7.3 Trajectory analysis

Production trajectories were unwrapped across periodic boundaries and stripped of solvent and ions. Analyses used MDTraj 1.10.3 (17), taking the first frame of the equilibrated production trajectory as the reference ("native") structure for each system. Native contacts were defined as heavy-atom pairs separated by less than 0.45 nm in the reference structure and quantified with the Best–Hummer switching function (18) with $\beta = 50\ nm^{-1}$ and $\lambda = 1.8$. Two complementary fraction-of-native-contacts metrics were computed. Intra-chain Q considered only contacts within the minibinder chain separated by at least three residues in sequence and therefore reports preservation of the binder fold rather than of binding. Interface Q considered contacts between the receptor and binder chains, with no sequence-separation criterion applied; native interface contact counts ranged from 255 to 503 depending on binder size. Binder Cα RMSD was calculated relative to the reference structure, and the change in radius of gyration reported as $\Delta Rg = Rg(t) - Rg(0)$.

##### 7.4 MM-GBSA binding free-energy calculations

Binding free energies were estimated using the single-trajectory MM-GBSA protocol implemented in MMPBSA.py (AmberTools; MMPBSA.py 14.0, sander 22.0) (19). Water-stripped trajectories were converted to Amber topologies with ParmEd 4.3.1 (20), mbondi2 generalized Born radii were assigned, and periodic boundary information was removed (IFBOX = 0) as required for implicit-solvent calculations. The first 10 ns of each production trajectory were discarded as additional equilibration and 500 evenly spaced frames extracted from the remainder for energy evaluation, using the modified generalized Born model at an ionic strength of 0.15 M. Per-residue energy decomposition was performed for interface residues, defined as residues within 6 Å of the partner chain. Reported binding free energies correspond to the DELTA TOTAL term.

##### 7.5 Statistical treatment

For every observable, each replicate trajectory was summarized by its time-averaged value, and the three replicates were then averaged to give a single value per design. Group comparisons between potent and non-potent designs used the design as the statistical unit (n = 7 and n = 12) and were evaluated by two-sided Mann–Whitney U tests, with Benjamini–Hochberg correction applied across the reported observables. Group summaries are shown in Figure S7 as medians with 95% bootstrap confidence intervals.

**Supplementary Data**

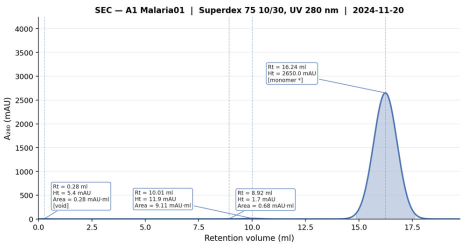

* Height estimated from axis scale; main peak table row scrolled off-screen in photo.

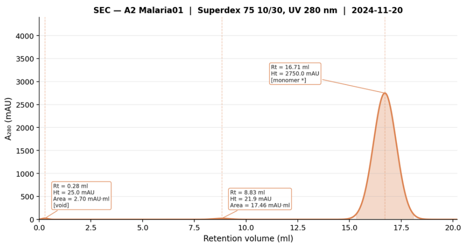

* Height estimated from axis scale; main peak table row scrolled off-screen in photo.

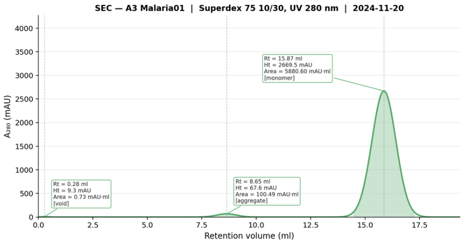

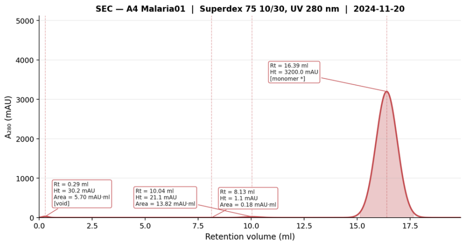

* Height estimated from axis scale; main peak table row scrolled off-screen in photo.

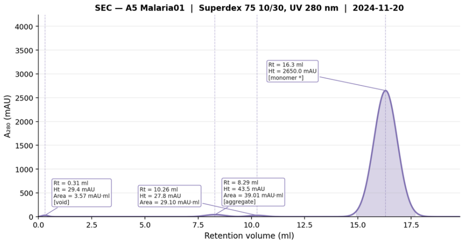

* Height estimated from axis scale; main peak table row scrolled off-screen in photo.

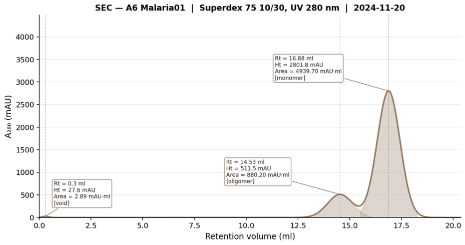

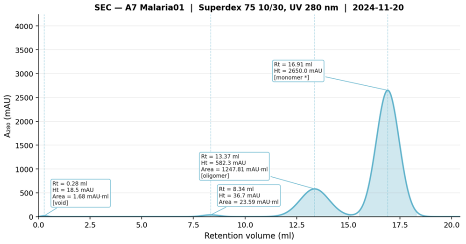

* Height estimated from axis scale; main peak table row scrolled off-screen in photo.

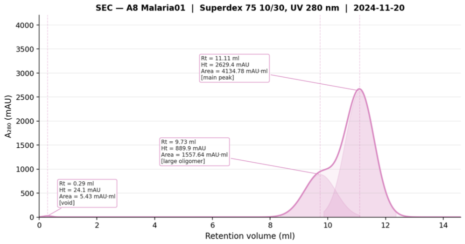

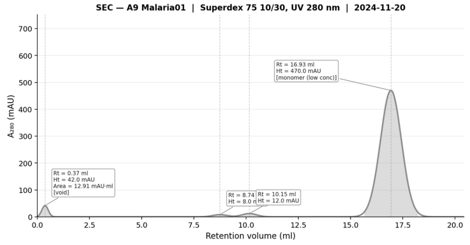

* Height estimated from axis scale; main peak table row scrolled off-screen in photo.

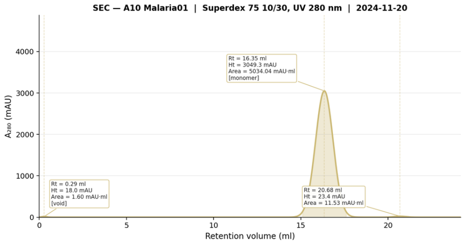

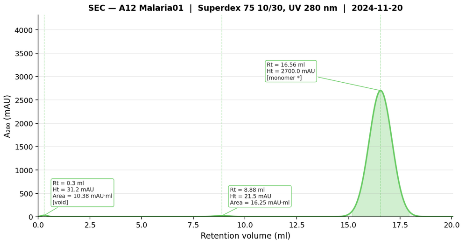

* Height estimated from axis scale; main peak table row scrolled off-screen in photo.

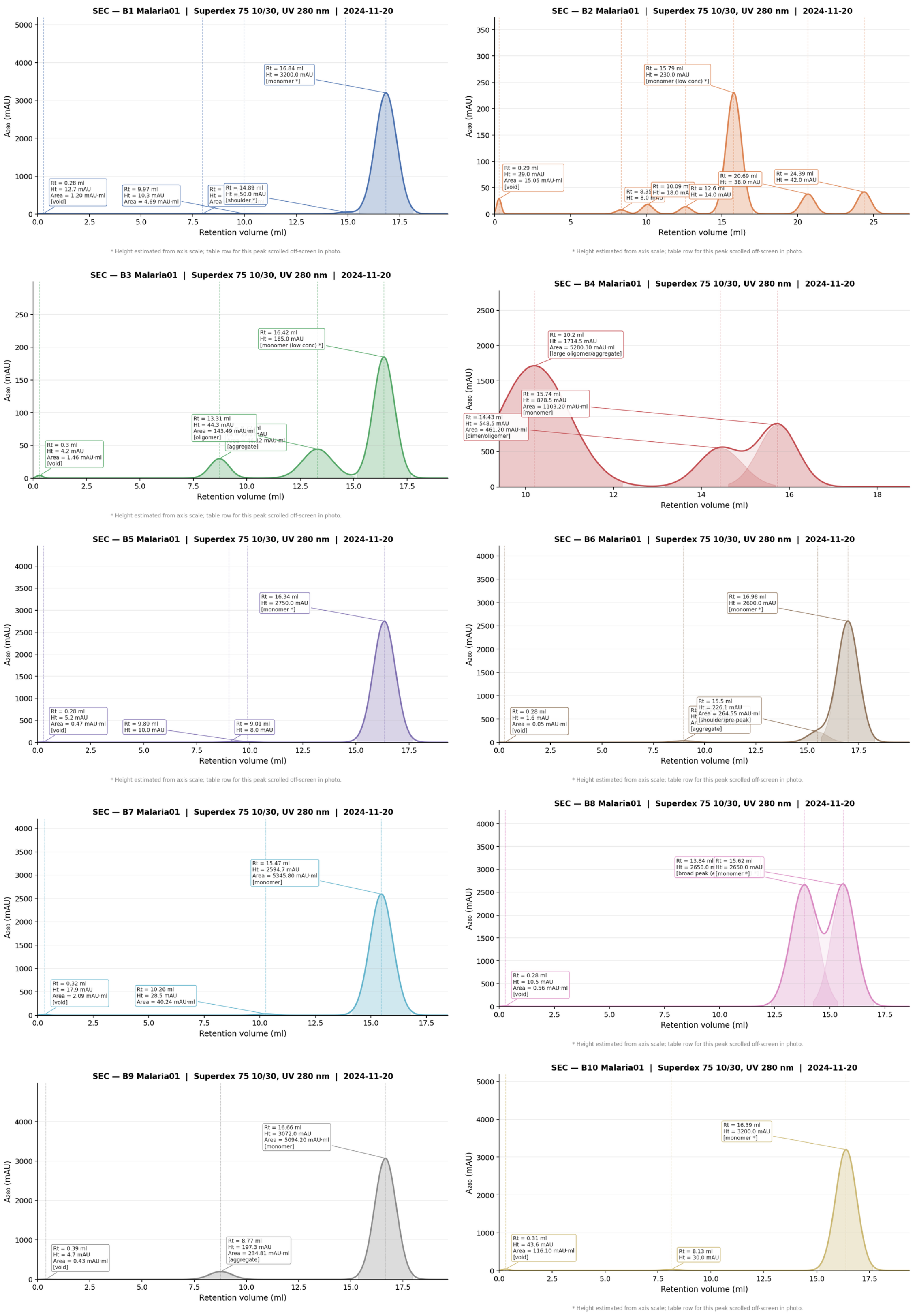

**Figure S1. Size-exclusion chromatography (SEC) of the designed minibinders.**

Analytical SEC traces (Superdex 75 10/300 GL; $A_{280}$) for 21 of the 23 designs, shown across two panels (mb-1 to mb-12 first, mb-13 to mb-24 second; mb-11 and mb-23 not available). Most designs displayed a dominant monomer peak; a subset (mb-6, mb-7, mb-8, mb-14, mb-15, mb-16, mb-20) showed additional higher-order (oligomer/aggregate) species. For every design carried forward, the monomeric peak fraction was resolved by SEC and pooled for downstream biophysical and functional characterization.

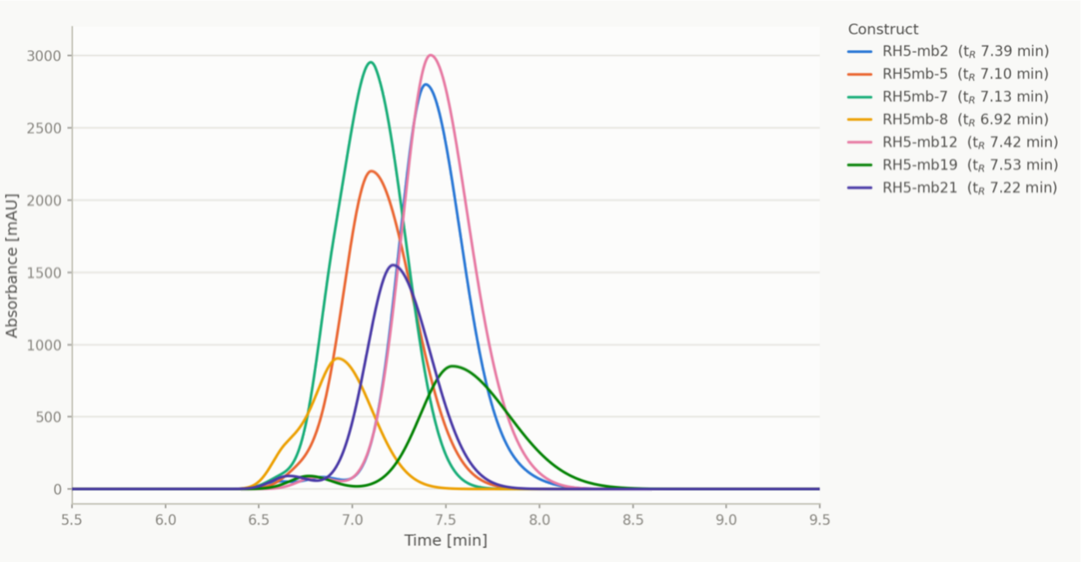

**Figure S2. Overlaid analytical HPLC-UV traces for seven RH5 minibinder constructs.**
RH5-mb2, RH5mb-5, RH5mb-7, RH5mb-8, RH5-mb12, RH5-mb19 and RH5-mb21 were run on HPLC column Biozen 3.0µm dSEC-1 90°A. All constructs elute as a dominant peak between 6.9 and 7.5 min. Traces are colored by construct identity. Curves were reconstructed from the peak apex times/heights annotated on the individual exported chromatogram images

**Table S1. AlphaFold3 validation metrics for representative designs.**

| Design | AF3 ipTM | AF3 pTM | Ranking score | Min PAE |
|---|---|---|---|---|
| RH5mb-8 | 0.91 | 0.68 | 1.02 | 1.25 |
| RH5mb-19 | 0.92 | 0.69 | 1.03 | 1.18 |
| RH5mb-21 | 0.89 | 0.67 | 0.99 | 1.32 |
| RH5mb-3 | 0.92 | 0.67 | 1.03 | 1.20 |

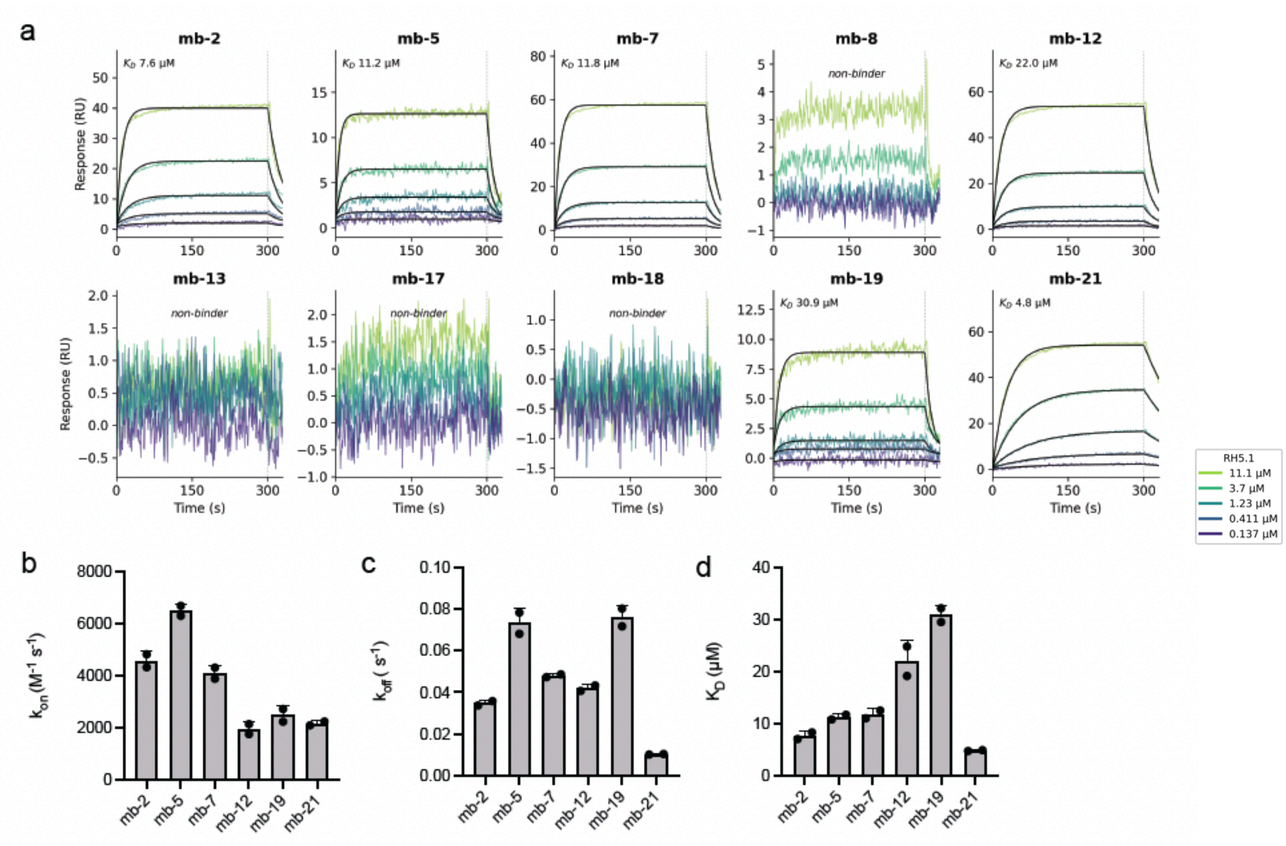

**Figure S3. SPR sensorgrams for all tested minibinders.**

(**a**) Reference-subtracted SPR sensorgrams (representative of two technical replicates) for all ten minibinders assayed by SPR (five-point, three-fold RH5.1 dilution series, 11.1–0.137 μM; the 33.3 μM injection was excluded for secondary-binding artefacts). Black, global 1:1 Langmuir fit to the association phase plus the first 30 s of dissociation (dissociation was biphasic over entire dissociation phase for some minibinders). The six binders (mb-2, mb-5, mb-7, mb-12, mb-19, mb-21) yield apparent $K_D$ values as indicated; mb-8, mb-13, mb-17 and mb-18 showed no measurable RH5 binding (non-binders). The measured $k_{on}$, $k_{off}$, and apparent $K_D$ for minibinders that bound RH5 in SPR are graphed in (**b**), (**c**), and (**d**), respectively.

**Table S2. SPR and BLI kinetic measurements.**

*Attached Excel Spreadsheet.*

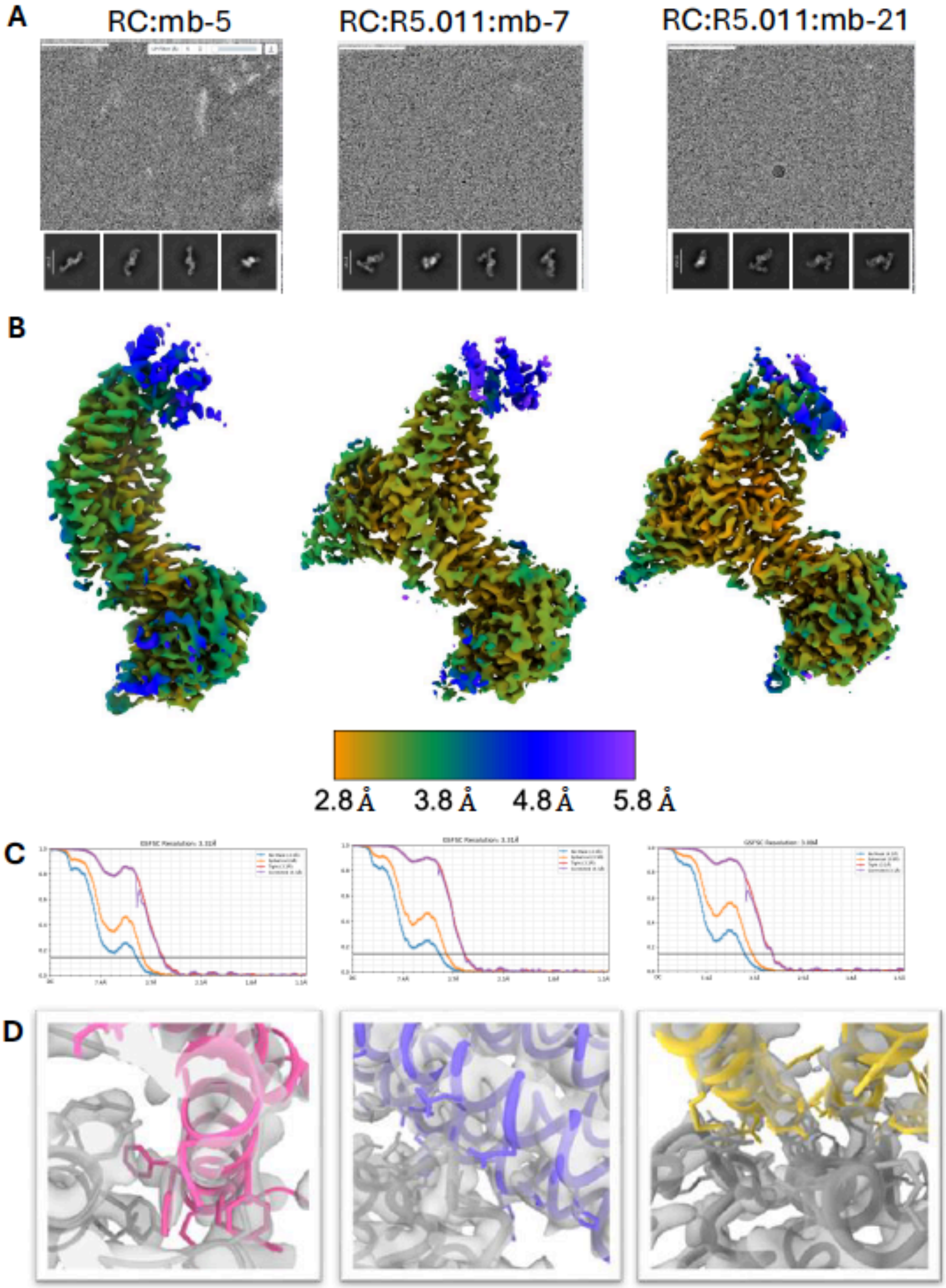

**Figure S4. Supporting Cryo-EM map data for minibinder complexes.**
(**a**) Representive micrographs and 2D classes for minbinder containing complexes (RH5:CyRPA:mb5, RH5:CyRPA:R5.011:mb-7, RH5:CyRPA:R5.011:mb-21). (**b**) Locally refined maps of the minibinder-containing complexes showing local resolution. The scale bar is shared between all maps. (**c**) Fourier-shell coefficient curves of each minibinder-complex locally refined map used to calculate the global resolution. (**d**) Examples of the Cryo-EM density at the minibinder:RH5 interface shown the EMready2 postprocessed maps. Mb-5 is shown in pink, mb-7 in purple and mb-21 in yellow. For mb-5 and mb-7, many of the bulkier and some smaller hydrophilic side chains

are resolved in the core of the minibinder-RH5 interface. For mb-7, Cryo-EM density for only a few bulky side chains is interpretable.

**Table S3. Cryo-EM data collection, refinement and validation statistics**

| | mb-5 bound to the RH5-CyRPA complex | mb-7 bound to the RH5-CyRPA-R5.011 complex | mb-21 bound to the RH5-CyRPA-R5.011 complex |
|---|---|---|---|
| **Data collection and processing** | | | |
| Magnification | 190000x | 190000x | 190000x |
| Voltage (kV) | 200 | 200 | 200 |
| Electron exposure (e–/Å$^2$) | 45 | 45 | 45 |
| Defocus range (μm) | -0.8 to -2.0 | -0.8 to -2.0 | -0.8 to -2.0 |
| Pixel size (Å) | 0.718 | 0.718 | 0.718 |
| Symmetry imposed | C1 | C1 | C1 |
| Final particle images (no.) | 91378 | 116699 | 192775 |
| Map resolution (Å) | 3.32 | 3.31 | 3.09 |
| FSC threshold | 0.143 | 0.143 | 0.143 |
| **Refinement** | | | |
| Initial model used (PDB code) | NA | NA | NA |
| Model resolution (Å) | 3.2/3.5 | 3.1/3.5 | 3.0/3.5 |
| FSC threshold | 0.143/0.5 | 0.143/0.5 | 0.143/0.5 |
| Map sharpening *B* factor (Å$^2$) | | | |
| Map Correlation Coefficient (Mask) | 0.79 | 0.8 | 0.72 |
| EMRinger Score | 1.93 | 3.1 | 2.75 |
| Model composition | | | |
| Non-hydrogen atoms | 6551 | 8425 | 8385 |
| Protein residues | 780 | 1021 | 1021 |
| Ligands | 0 | 0 | 0 |
| *B* factors (Å$^2$) | 71.8 | 75.2 | 69.7 |
| Protein | 57.37 | 54.47 | 54.69 |
| Ligand | NA | NA | NA |

| | | | |
|---|---|---|---|
| R.m.s. deviations | | | |
| Bond lengths (Å) | 0.008 | 0.003 | 0.004 |
| Bond angles (°) | 0.856 | 0.725 | 1.101 |
| Validation | | | |
| MolProbity score | 1.89 | 1.43 | 1.51 |
| Clashscore | 8.32 | 7.83 | 9.67 |
| Poor rotamers (%) | 0.82 | 0 | 0 |
| Cβ outliers | NA | NA | NA |
| Ramachandran plot | | | |
| Favored (%) | 93.26 | 99.5 | 98.32 |
| Allowed (%) | 6.74 | 0.3 | 0.99 |
| Disallowed (%) | 0 | 0.2 | 0.69 |

**Table S4. PDBePISA analysis of minibinder interfaces (21).**

*Attached Excel Spreadsheet.*

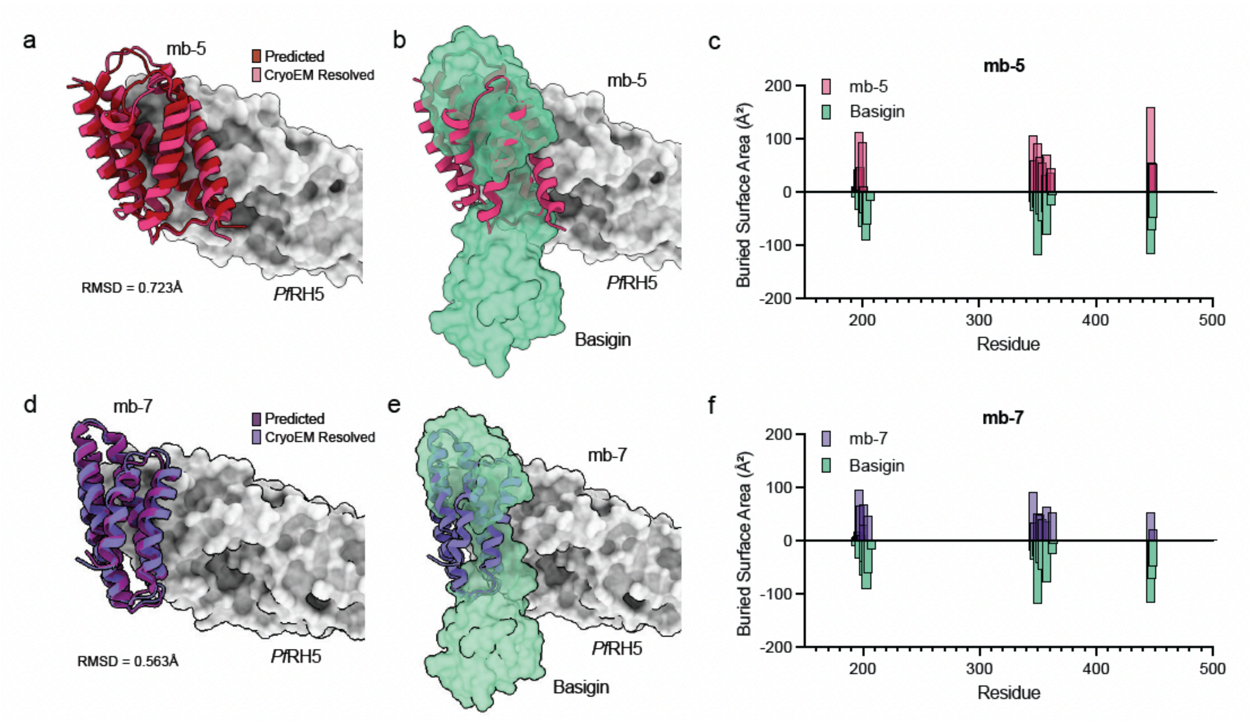

#### Figure S5. CryoEM structures of mb-5, mb-7 minibinders complexed with *Pf*RH5.

(**a, d**) Experimental CryoEM structure of mb-5 and mb-7 complexed with *Pf*RH5 (grey). Predicted mb-5 (dark red) and mb-7 (dark purple) are overlaid. (**b, e**) The same mb-5 and mb-7 minibinders and *Pf*RH5 complexes superposed with the erythrocyte receptor basigin (green surface). (**f**) Per-residue buried surface area (BSA) on *Pf*RH5 for mb-5, mb-7 and for basigin. The minibinders and basigin bury an overlapping set of RH5 residues, clustered around ~195–205, ~345–370, and ~440–450, the same hotspots basigin engages.

#### Table S5. Model-to-experiment structural comparison for CryoEM-resolved minibinders.

| Minibinder | *Dd2* $IC_{50}$ | Exp. / pred. RH5 footprint | Footprint Jaccard | Basigin coverage | Hotspots | Angle vs prediction (°) | Experimental total BSA (Å²) |
|---|---|---|---|---|---|---|---|
| RH5mb-5 | 107 nM | 23 / 22 | 0.800 | 0.789 | 3 | 3.39 | 2219 |
| RH5mb-7 | 201 nM | 20 / 20 | 0.905 | 0.789 | 3 | 5.60 | 1606 |
| RH5mb-21 | 193 nM | 28 / 29 | 0.727 | 0.737 | 2 | 3.35 | 2705 |

*Footprints and basigin coverage use the common 5Å heavy-atom contact definition. Total BSA is the summed Shrake-Rupley loss from both binding partners.*

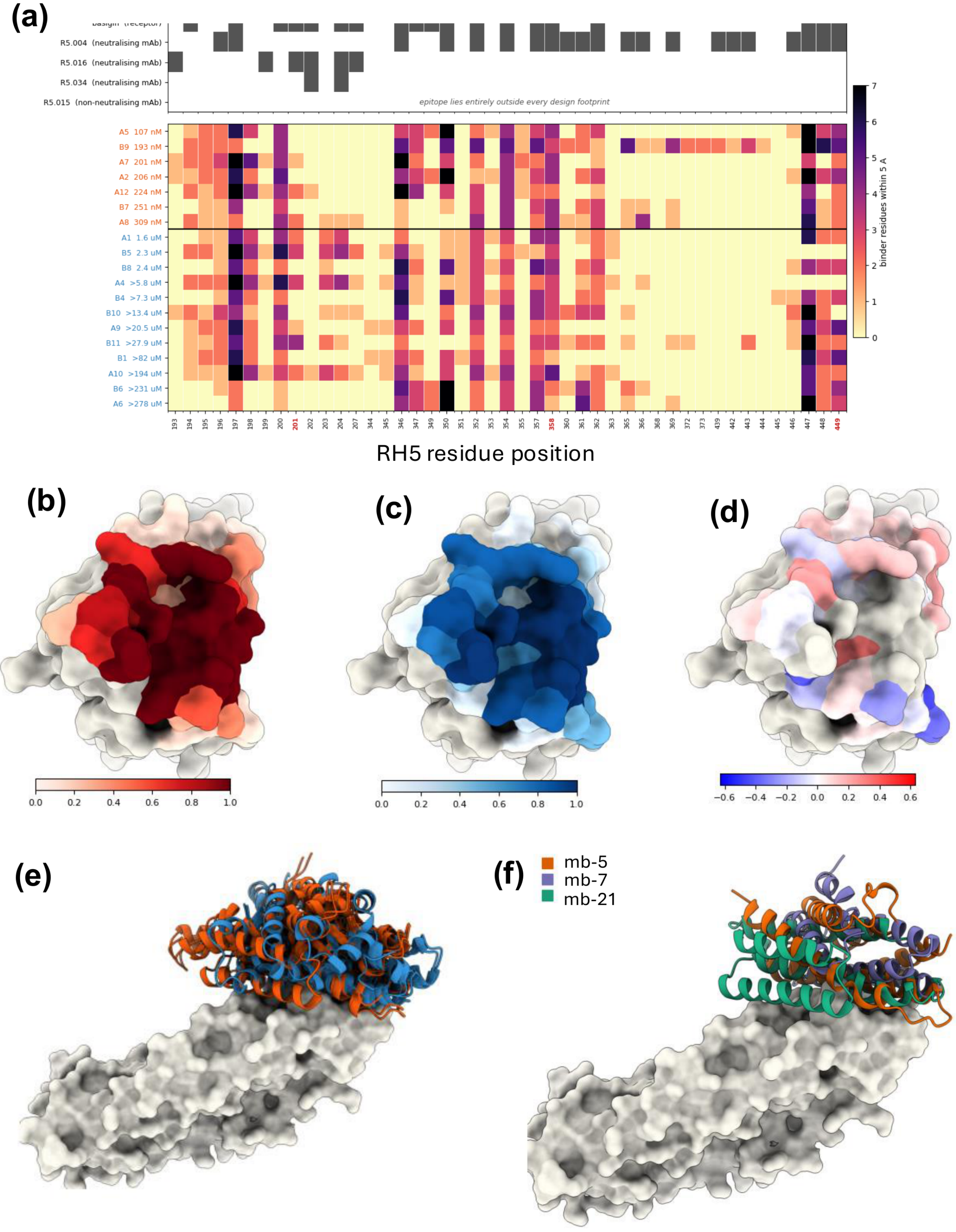

**Figure S6. Designed minibinders engage a common RH5 surface irrespective of potency.**

(a) RH5 contact fingerprints at design-sampled positions (5 Å heavy-atom cutoff). Reference complexes are shown above; the 19 designs below, ordered by potency and separated by a horizontal rule into potent and non-potent groups. Color indicates the number of minibinder residues within 5 Å of each RH5 position. Seven positions (197, 200, 346, 352, 354, 357 and 358) are contacted by all 19 designs, and the epitope of the non-neutralizing antibody R5.015 lies entirely outside the design-sampled region. (b–d) RH5 surface occupancy for potent designs, non-potent designs, and the potent-minus-non-potent difference, colored by the fraction of designs in each group contacting every residue. (e) Side-on overlay of the predicted minibinder ensembles for both groups after RH5 registration. (f) Experimental mb-5, mb-7 and mb-21 cryo-EM minibinders shown in the same common RH5 reference frame (chain A of PDB 4WAT). Each experimental lead closely reproduces its own predicted footprint (Jaccard 0.800, 0.905 and 0.727, respectively) and coarse approach direction (3.39°, 5.60° and 3.35° from prediction), while the three leads differ substantially from one another (pairwise footprint Jaccard 0.72, 0.55 and 0.45). Quantitative comparisons are given in Table S5.

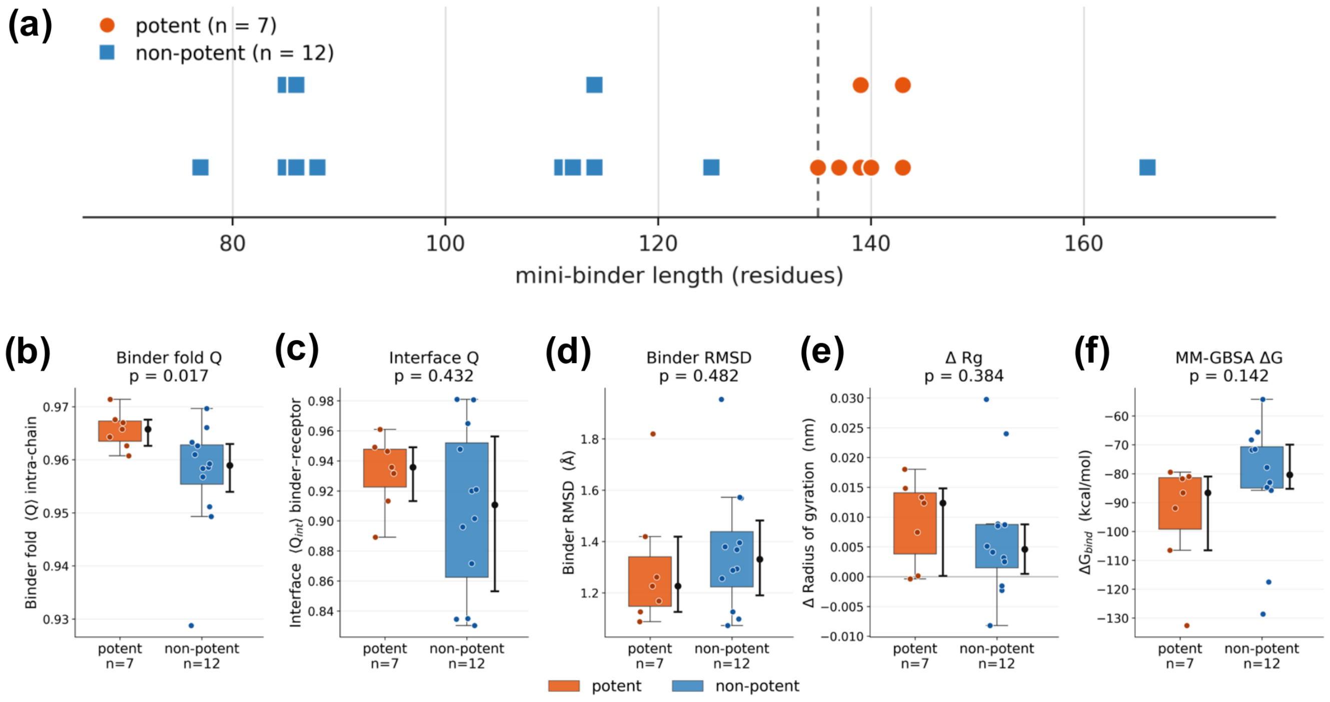

**Figure S7. Potency is not resolvable from computational features.**

(a) Minibinder length for the 19 modelled designs, by potency group. Points sharing a length are stacked vertically (vertical position carries no meaning). Dashed line, 135 residues. All seven potent designs fall within a narrow 135–143 residue window, whereas eleven of twelve non-potent designs are shorter (77–125 residues). The seven potent designs derive from six distinct scaffolds, so the length window does not reflect a shared lineage. (b–f) Molecular dynamics and MM-GBSA properties of the predicted complexes, simulated in triplicate for 100 ns each (7.2 μs aggregate production sampling). Each point is one minibinder, plotted as the mean of its three independent replicates; boxes show the median and interquartile range, with the adjacent bar giving the median and 95% bootstrap confidence interval. (b) Intra-chain Q, reporting stability of the minibinder fold. (c) Interface Q, the fraction of native receptor–binder contacts retained and the binding-relevant measure. (d) Binder Cα RMSD. (e) Change in radius of gyration. (f) MM-GBSA binding free energy. No binding-relevant metric differed between groups (Q_int $p = 0.43$; RMSD $p = 0.48$; ΔRg $p = 0.38$; ΔG $p = 0.14$), and no comparison reached significance after Benjamini–Hochberg correction across the five observables (minimum $q = 0.085$). The only nominally significant contrast was intra-chain Q ($p = 0.017$), which reports the binder's own fold rather than the interface. Geometry comparisons are reported in Table S7.

**Table S6. $IC_{50}$ values of all tested minibinders.**

$IC_{50}$ values for biotinylated minibinders used for SPR (AviTag-) are also included. Biotinylated minibinders demonstrate similar potency patterns as non-biotinylated minibinders (Figure S8).

*Attached Excel Spreadsheet.*

**Table S7. Selected structural geometry comparisons between potent and non-potent minibinders.**

| Metric | Potent median | Non-potent median | Cliff δ | Design exact p | Scaffold exact p | BH q |
|---|---|---|---|---|---|---|
| Basigin coverage | 0.789 | 0.737 | +0.381 | 0.047 | 0.442 | 0.639 |
| Total buried SASA (Å²) | 2180 | 1990 | +0.405 | 0.166 | 0.321 | 0.639 |
| Buried SASA / binder residue (Å²) | 16.1 | 20.9 | -0.619 | 0.224 | 0.323 | 0.639 |
| Binder standoff (Å) | 15.1 | 13.5 | +0.238 | 0.228 | 0.590 | 0.639 |
| Footprint radius of gyration (Å) | 9.19 | 10.2 | -0.095 | 0.306 | 0.701 | 0.710 |
| Residue-contact pairs | 58 | 55.5 | +0.262 | 0.533 | 0.127 | 0.830 |
| RH5 footprint residues | 22 | 21 | +0.060 | 0.953 | 0.776 | 1.000 |
| Maximum neutralizing-mAb Jaccard | 0.483 | 0.475 | +0.155 | 0.989 | 0.501 | 1.000 |
| Designed hotspots contacted | 2 | 2 | +0.143 | 1.000 | 1.000 | 1.000 |

*BH q values refer to the design-level geometry family. No conventional interface-size, contact-count, or epitope-overlap metric remains significant after multiple-testing correction. Basigin coverage is nominal at the design level but is not supported at the scaffold level.*

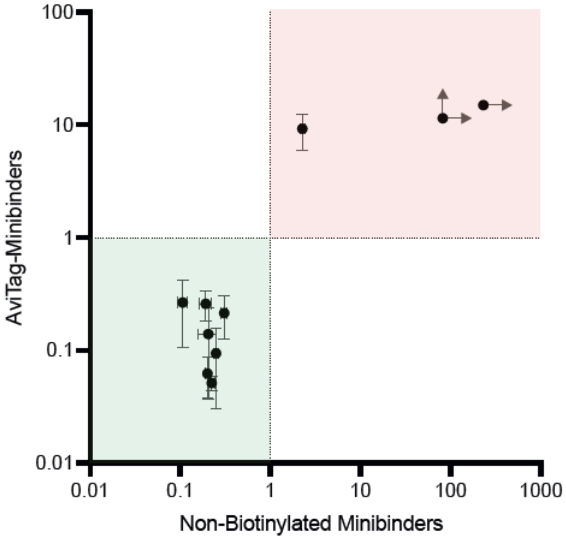

**Figure S8. Comparison of biotinylated (AviTag-) vs. non-biotinylated minibinder $IC_{50}$ values.** To verify that biotinylated minibinders used for kinetics assays via SPR exhibit similar potency patterns with non-biotinylated minibinders used for potency assays, we compared their $IC_{50}$ values. All potent minibinders (<1μM; green shading) remain potent regardless of the presence/absence of biotinylation. Similarly, all non-potent minibinders remain non-potent (red shading) regardless of biotinylation state. Each point represents an individual minibinder sequence, with error bars indicating standard error (Supplementary Table S6). Arrows on the least potent minibinders indicate undefined $IC_{50}$ values that would not be determined below the maximum tested concentrations.

**Table S8. Mutated amino acids in field-relevant *P. falciparum* isolate *Pf*RH5 relative to *Dd2* reference.**

| Strain | Amino Acid Position | Reference Amino Acid | Mutated Amino Acid |
|---|---|---|---|
| Mozambique (R13.24-B) | 147 | Y | H |
| Mozambique (R13.24-B) | 148 | H | D |
| Mozambique (R13.24-B) | 203 | C | Y |
| Uganda (MAS156) | 203 | C | Y |

**Table S9. Potent *Dd2*-Luc RH5 minibinders retain sub-micromolar potency against field-relevant *P. falciparum* isolates.**

IC$_{50}$ values (µM; arithmetic mean ± SE, with number of biological replicates in parentheses) against the laboratory strain *Dd2*-Luc and two field isolates, Uganda (MAS156) and Mozambique (R13.24-B). n.d., not determined; “>” denotes IC$_{50}$ above the highest concentration tested (inactive).

| Compound/Molecule | *Dd2*-Luc | Uganda (MAS156) | Mozambique (R13.24-B) |
|---|---|---|---|
| **Reference controls** | | | |
| **GNF-179** | 0.000817 ± 0.00019 (7) | 0.00179 ± 1.5e-05 (2) | 0.00448 ± 0.0014 (2) |
| **R5.004** | 0.0433 ± 0.0058 (6) | 0.0675 ± 0.0048 (2) | 0.0421 ± 0.016 (2) |
| **Potent minibinders (Dd2-Luc IC50 < 1 µM)** | | | |
| **mb-2** | 0.206 ± 0.046 (5) | 0.56 ± 0.0025 (2) | 0.323 ± 0.1 (2) |
| **mb-5** | 0.107 ± 0.013 (5) | 0.35 ± 0.0005 (2) | 0.166 ± 0.005 (2) |
| **mb-7** | 0.201 ± 0.0078 (5) | 0.293 ± 0.0015 (2) | 0.122 ± 0.0085 (2) |
| **mb-8** | 0.309 ± 0.021 (3) | n.d. | 0.83 ± 0.081 (2) |
| **mb-12** | 0.224 ± 0.017 (5) | 0.603 ± 0.0095 (2) | 0.213 ± 0.057 (2) |
| **mb-19** | 0.251 ± 0.012 (3) | 0.0361 (1) | 0.17 ± 0.0065 (2) |
| **mb-21** | 0.193 ± 0.031 (8) | 0.467 ± 0.056 (2) | 0.266 ± 0.095 (2) |
| ***Inactive / weak minibinders*** | | | |
| **mb-1** | 1.58 ± 0.071 (3) | n.d. | > 12 (2) |
| **mb-13** | > 82 (5) | > 12 (1) | > 12 (2) |
| **mb-17** | 2.27 ± 0.12 (3) | n.d. | > 12 (2) |

**Table S10. Parasitemia data and statistical comparisons of stage progression assays.**

*Attached Excel spreadsheet.*

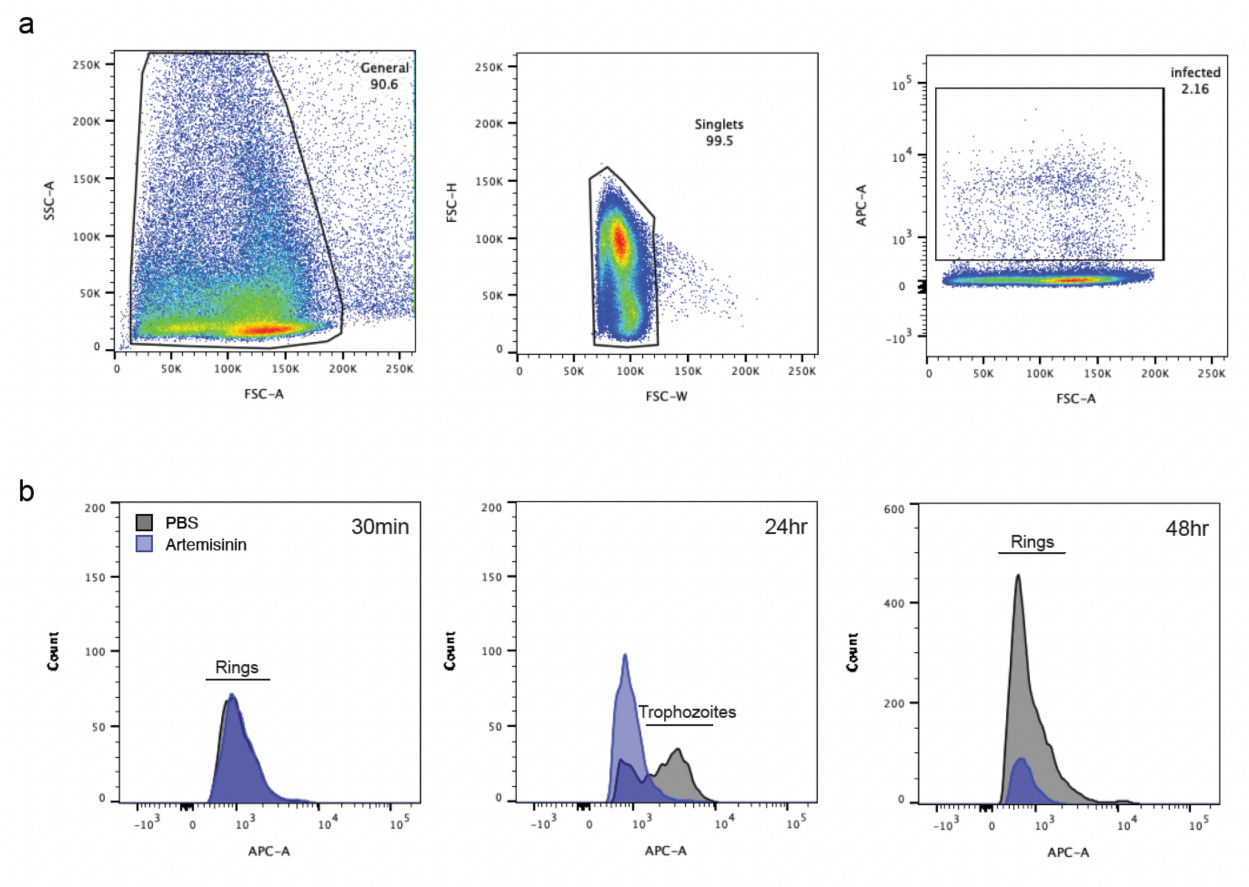

**Figure S9. Representative gating strategy to identify parasite-infected erythrocytes and stage.** (**a**) Initial general gating via size/granularity (FSC-A, SSC-A), before exclusion of doublets via FSC-W vs. FSC-H. Infected erythrocytes are identified via APC positive events above uninfected threshold. (**b**) Stage of parasite at each respective time point in stage-progression assays via histogram of APC-A from the infected gating population of (**a**). Artemisinin and PBS were used as negative and positive controls, respectively, to compare against for all other conditions to analyze stage-specificity of treatment. Artemisinin, an all-stage antimalarial, inhibits trophozoite formation and thus decreases the proportion of successful reinvasion events relative to PBS control. Rings are identified as lower APC signal events relative to higher APC signal in trophozoites, reflecting relative DNA content at their respective stages.

### Cited Literature
